# Slide-seq: A Scalable Technology for Measuring Genome-Wide Expression at High Spatial Resolution

**DOI:** 10.1101/563395

**Authors:** Samuel G. Rodriques, Robert R. Stickels, Aleksandrina Goeva, Carly A. Martin, Evan Murray, Charles R. Vanderburg, Joshua Welch, Linlin M. Chen, Fei Chen, Evan Z. Macosko

**Author notes:** These authors contributed equally to this work (Samuel G. Rodriques, Robert R. Stickels). These authors contributed equally to this work (Fei Chen, Evan Z. Macosko).

## Abstract

The spatial organization of cells in tissue has a profound influence on their function, yet a high-throughput, genome-wide readout of gene expression with cellular resolution is lacking. Here, we introduce Slide-seq, a highly scalable method that enables facile generation of large volumes of unbiased spatial transcriptomes with 10 µm spatial resolution, comparable to the size of individual cells. In Slide-seq, RNA is transferred from freshly frozen tissue sections onto a surface covered in DNA-barcoded beads with known positions, allowing the spatial locations of the RNA to be inferred by sequencing. To demonstrate Slide-seq’s utility, we localized cell types identified by large-scale scRNA-seq datasets within the cerebellum and hippocampus. We next systematically characterized spatial gene expression patterns in the Purkinje layer of mouse cerebellum, identifying new axes of variation across Purkinje cell compartments. Finally, we used Slide-seq to define the temporal evolution of cell-type-specific responses in a mouse model of traumatic brain injury. Slide-seq will accelerate biological discovery by enabling routine, high-resolution spatial mapping of gene expression.

**One Sentence Summary:** Slide-seq measures genome-wide expression in complex tissues at 10-micron resolution.

## Main Text

The functions of complex tissues are fundamentally tied to the organization of their resident cell types. However, unbiased methods for exploring, genome-wide, spatial distributions of gene expression in tissues are lacking. Recently developed multiplexed *in situ* hybridization and sequencing-based approaches measure gene expression within cells and tissues with subcellular spatial resolution (*1*, *2*), but can be laborious, and require specialized knowledge and equipment. In addition, most *in situ* approaches require the upfront identification and selection of specific target genes for measurement, which may limit *de novo* discovery of spatially varying genes. By contrast, previous technologies for spatially encoded RNA-sequencing using barcoded oligonucleotide capture arrays are presently limited to resolutions in the hundreds of microns (*3*), which is insufficient for detecting many important tissue features.

To develop Slide-seq, we first asked whether barcoded oligonucleotides could be arrayed randomly on a surface at high spatial resolution, with locations determined post hoc. We packed uniquely barcoded 10 µm microparticles (‘beads’)—similar to those used by the Drop-seq approach to scRNA-seq (*4*)—onto a rubber-coated glass coverslip forming a monolayer we termed a “puck” (**Fig. S1**, 92.9% ± 2.15% packing). We found that the bead barcode sequences on the puck could be uniquely determined via SOLiD sequencing-by-ligation chemistry (Fig. 1A, **Fig. S1**). Moreover, pucks could be stored for extended periods of time prior to use, allowing them to be produced in large batches and used as needed.

**Figure 1:**
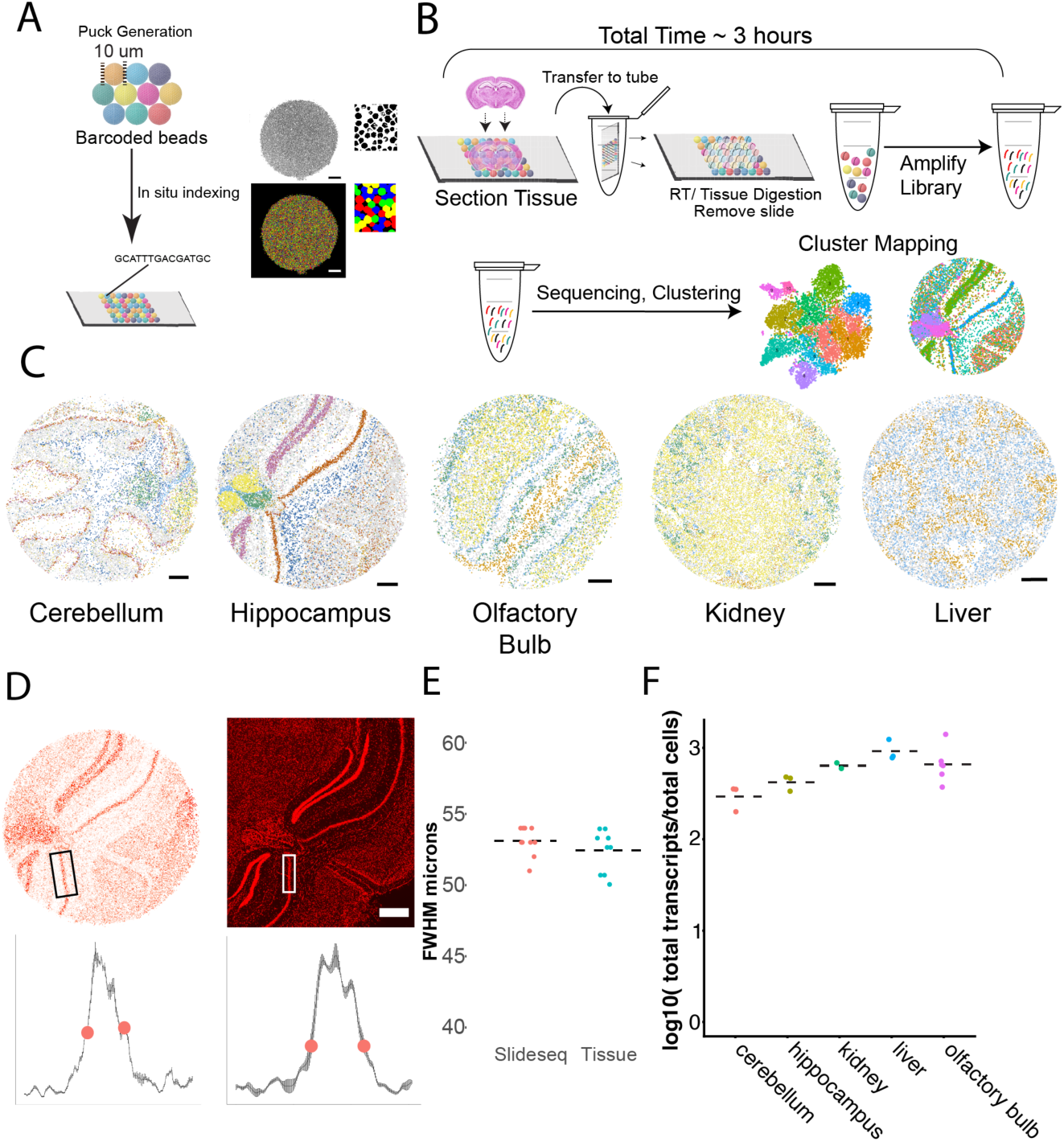
High-resolution RNA capture from tissue by Slide-seq. (**A**) Left: Schematic of array generation. A monolayer of randomly deposited, DNA barcoded beads (termed a “puck”) is spatially indexed by SOLiD sequencing. Top Right: A representative puck with called barcodes shown in black. Bottom Right: A composite image of the same puck colored by the base calls for a single base of SOLiD sequencing. (Scale bar 500 μm) (**B**) Top Row: Schematic of the sample preparation procedure developed for Slide-seq. Total time for library generation is ~3 hrs. Bottom Row: Schematic of a naïve analysis, in which each bead is clustered by its gene expression, visualized in a tSNE two-dimensional embedding, and by locations in space. (**C**) Spatial positions of Slide-seq beads, colored by clusters defined purely by gene expression relationships amongst beads, across five tissue types (see **Fig. S2** for tSNE embeddings and definitions). (**D**) Characterization of lateral diffusion of signal on the Slide-seq surface. Top Left: Digital image of a Slide-seq puck with bead color intensity scaled by total transcript counts. Top Right: Image of the adjacent tissue section, stained with DAPI (scale bar 500 μm). Boxes represent regions where an intensity profile was taken across CA1. Bottom left: Profile of pixel intensity across CA1 in Slide-seq. Bottom right: Profile across CA1 in DAPI stained tissue. Red dots represent locations of half max of the distribution. (**E**) Quantification of full width at half maximum of profiles in (D), from both Slide-seq (red dots) and DAPI-stained tissue (blue dots) (dotted line, mean; N = 10 profiles) (**F**) Log ratio of total number of quantified RNA transcripts on a puck to the number of cells counted on a serially stained DAPI slice of equal area (dotted line, mean) across five different tissues.

Next, to capture RNA from tissue with high resolution, we developed a protocol wherein fresh-frozen tissue sections (10 µm thickness) were transferred onto the dried bead surface via cryosectioning (*4*). We observed efficient hybridization of tissue-extracted mRNA to polyT capture sequences, after which 3’-end digital expression libraries could be prepared and sequenced (*5*). The simplicity of the protocol enabled fast, facile generation of libraries in a highly multiplexed fashion (Fig. 1B). To highlight its generalizability, we performed Slide-seq across a range of samples, generating data from three different organs (mouse brain, kidney, liver). Expression measurements by Slide-seq agreed well with those from bulk mRNAseq (r = 0.89) (**Fig. S1D**), similar to standard single-cell profiling technologies (*5*), and average mRNA transcript capture per cell was consistent across tissues and experiments (Fig. 1F).

Unbiased clustering of individual bead profiles using single-cell analysis approaches (*4*) yielded cluster assignments reflecting known positions of cell types in the assayed tissues (Fig 1C). Specifically, in analyses of three different regions of brain—cerebellum, hippocampus, and olfactory bulb—the different neuronal cell types that form the layered tissue architecture were immediately detectable, as were populations of resident glial types. In kidney, we identified clusters of cells representing podocytes, surrounded by proximal and distal tubule populations. In liver, the lobule architecture was visible, with genes known to manifest zonation patterns driving the distinction between two main clusters (*6*). To quantify diffusion in the tissue, we compared the width of mRNA transcript density in hippocampal CA1 observed in Slide-seq to that observed in an adjacent, DAPI-stained tissue section (Fig. 1D). We estimated the length-scale of lateral diffusion of transcripts during hybridization to be 1.4um ± 1.3um (Fig. 1E), implying that mRNA is transferred from the tissue to the beads with high spatial resolution.

Positioning cell types defined by large-scale scRNA-seq datasets represents an important application of spatial technologies. To map scRNA-seq cell types onto Slide-seq data, we developed a computational approach called Non-negative Matrix Factorization Regression (NMFreg). NMFreg reconstructs expression of each Slide-seq bead as a weighted combination of metagene factors, each corresponding to the expression signature of an individual cell type, defined from scRNA-seq (Fig. 2A). Application of NMFreg to a Slide-seq puck yields a quantitative estimate of the contriubution of each scRNA-seq-defined cell type to each. Bead, allowing the locations of cell types in space to be inferred. Using cell type definitions from a recently acquired scRNA-seq dataset of the mouse cerebellum (*7*), NMFreg on a mouse cerebellar puck recapitulated the spatial distributions of classical neuronal and non-neuronal cell types, such as granule cells, Golgi interneurons, unipolar brush cells, Purkinje cells, and oligodendrocytes (Fig. 2B). We found that 65.8% +/-1.4% of beads showed significant representation (defined by at least 25% loading) of only one cell type (*4*), whereas 32.6% +/-1.2% showed significant representation of two cell types (mean ± std, N=7 cerebellar pucks) (Fig. 2C **left, Fig. S3**). The high spatial resolution of the method was found to be key for assigning beads to individual cell types with high confidence: upon artificially aggregating the data into larger feature sizes, we failed to confidently map cell types in heterogeneous regions of tissue, while homogenous regions such as the granular layer of the cerebellum remained reasonably cell-type-specific (**Fig. S4**). Importantly, the representation of cell types in Slide-seq more accurately represented the spatial distribution of cell types expected from immunohistochemistry than single-cell sequencing, thus allowing for better identification of rare cell types with distinct spatial organization: whereas Purkinje neurons make up only 0.7% of cerebellar single-cell atlas data, they make up 7.8% ± 1.3% (mean ± std, N=7 pucks) of the area of a cerebellar puck (Fig. 2C, **right**).

**Figure 2:**
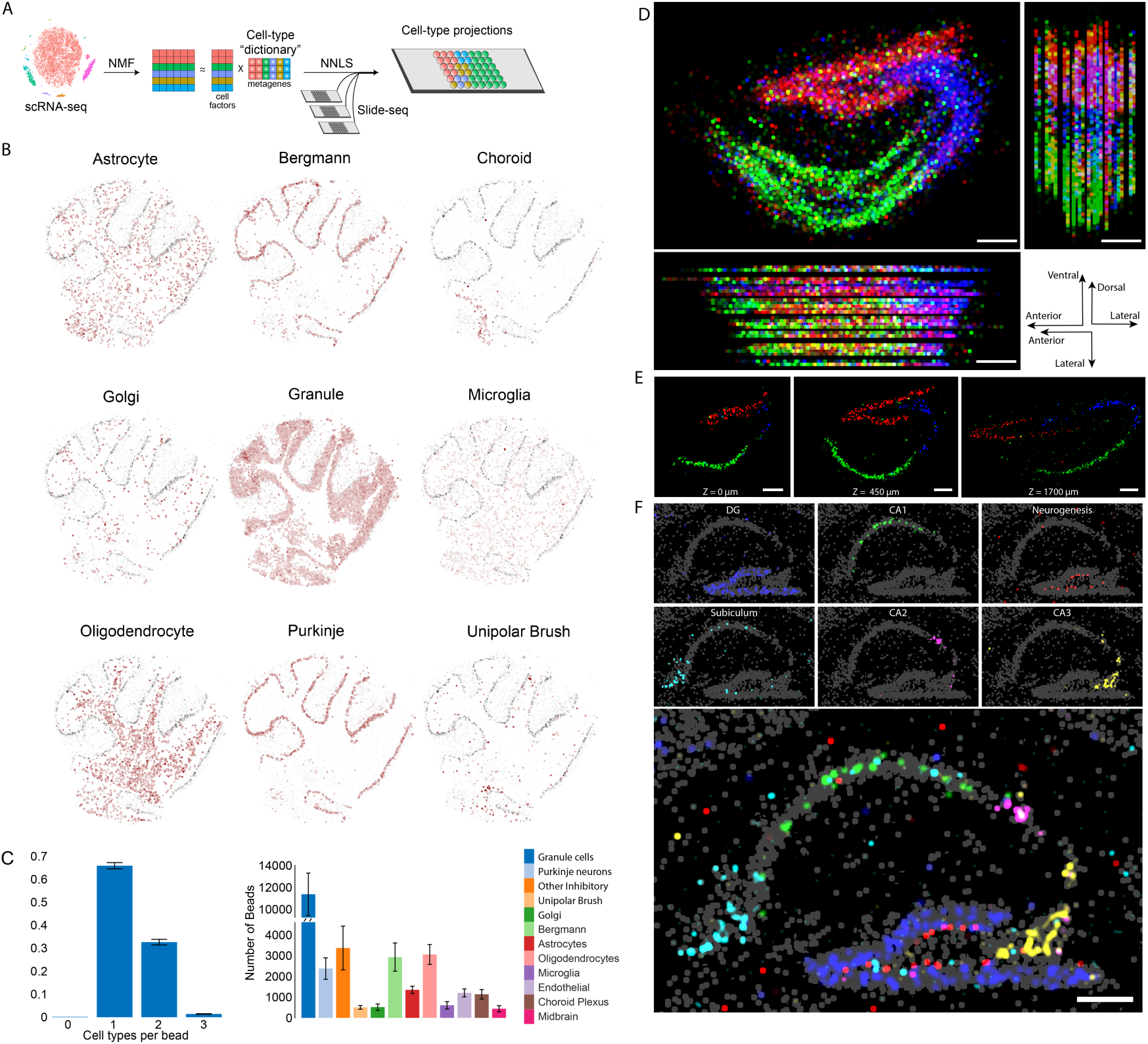
Localization of cell types in cerebellum and hippocampus using Slide-seq. (**A**) Schematic for assigning cell types from scRNA-seq datasets to Slide-seq beads using NMF and NNLS regression (NMFreg). (**B**) Loadings of individual cell types, defined by scRNA-seq cerebellum (*7*) on each bead (minimum 15 genes per bead) of one 3 mm-diameter coronal cerebellar puck (red, cell type location, gray, Purkinje loadings plotted as a counterstain). (**C**) Left: Number of cell types assigned per bead. A cell type was assigned to a bead if the factors corresponding to that cell type made up at least 25% of bead loading (**Fig. S3**). Right: The number of beads called as each atlas-defined cell type for cerebellar pucks. Error bars represent standard deviation (N=7 pucks). (**D**) Projection of hippocampal volume with NMFreg cell type calls for CA1 (green), CA2/3 (blue) and dentate gyrus (Red). Top left: Sagittal projection. Top right: Coronal projection. Bottom left: Horizontal projection. Bottom right: axis orientations for each of the projections. (**E**) Cell type calls of three representative sections from the dataset with the position on the mediolateral axis denoted at the bottom of the image. (**F**) Top: Metagene profiles on a sagittal hippocampus section representing cell subtypes. Bottom: Composite image of all metagenes. All scale bars show 250 µm.

To demonstrate the scalability of Slide-seq, we applied it to 66 tissue slices from a single dorsal mouse hippocampus, covering a volume of 39 cubic millimeters, with roughly 10 µm resolution in the dorsal-ventral and anterior-posterior axes, and ~20 µm resolution in medial-lateral axis. This region contained approximately 1 million beads that could be confidently assigned to single cell types. We computationally co-registered pucks along the medial-lateral axis, allowing for visualization of the cell types and gene expression in the hippocampus at high resolution in three dimensions (Fig. 2D, E, **Supplementary Video 1**). We plotted metagenes comprised of markers—defined by a recent large-scale single-cell study (*7*)—for the dentate gyrus, CA2, CA3, a subiculum subpopulation, an anteriorly localized CA1 subset (exemplified by the marker *Tenm3*) and cells undergoing mitosis and neurogenesis. The metagenes were highly expressed and specific for the expected regions (Fig. 2F), confirming the ability of Slide-seq to localize both common cell-types as well as subtler cellular subpopulations. The entire experimental processing for these 66 pucks (excluding puck generation) required roughly 40 person-hours (*4*), and only standard experimental apparatus associated with cryosectioning and next-generation sequencing, making Slide-seq readily scalable to measure gene expression in large tissue volumes.

One key advantage of Slide-seq’s high-resolution, genome-wide approach is the ability to identify spatially variable genes within specific cell types. To find genes with non-random spatial patterns, we developed a nonparametric, kernel-free algorithm to identify genes with spatially non-random distribution across the puck (**Fig. S5**) (*4*). Application of this algorithm to a coronally sliced cerebellum puck identified *Ogfrl1, Prkcd* and *Atp2b1* as possessing highly non-random patterns of expression localized just inferior to the cerebellum (Fig. 3A). We found *Ogfrl1* in particular to be a highly specific marker for PV interneurons in the molecular and fusiform layers of the dorsal cochlear nucleus (Fig. 3B), likely the cartwheel cells of the dorsal cochlear nucleus that are thought to be involved in the generation of feedforward inhibition (*8*, *9*). The identification of marker genes for this population may assist in future optogenetic and labeling studies of the cochlear nucleus.

**Figure 3:**
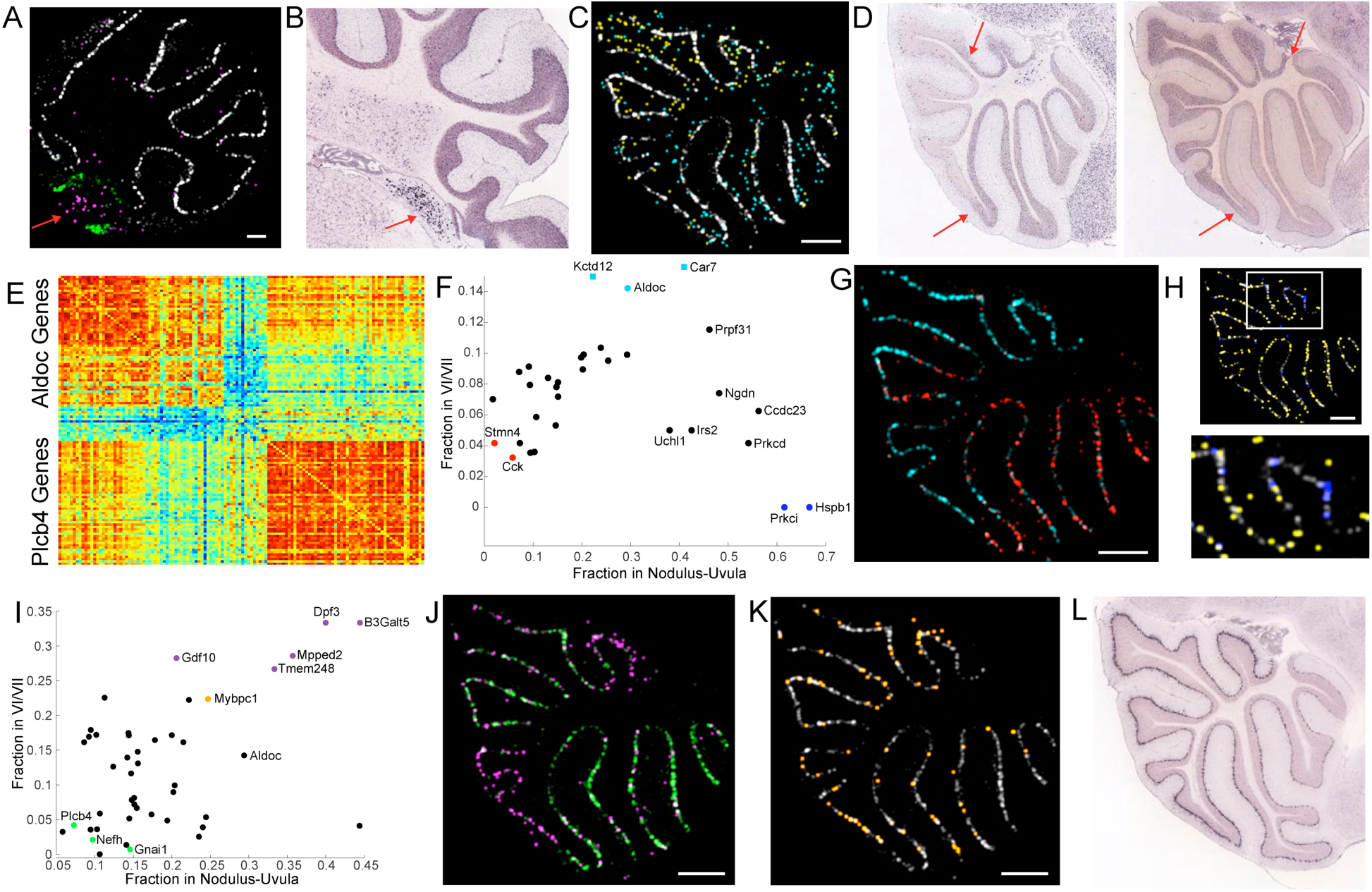
Identification of novel variation in cerebellar gene expression by Slide-seq. (**A**) A coronal cerebellar puck is shown, with Purkinje-assigned beads in white, choroid-assigned beads in green, and beads expressing *Ogfrl1* in magenta. Red arrow indicates cluster of *Ogfrl1*-positive beads. (**B**) An Allen ISH atlas image of *Ogfrl1*, from a similar brain region. Red arrow indicates *Ogfrl1* expression in the cochlear nucleus. (**C**) A sagittal cerebellar puck showing counts of *Pcp4* (gray), *Rasgrf1* (blue), and a metagene consisting of *Gprin3*, *Cemip*, *Mab21l2*, and *Syndig1l* (yellow). (**D**) Allen atlas images of *Rasgrf1* (left) and *Gprin3* (right). Arrows indicate point of boundary of expression within the granular layer for each gene. (**E**) A heatmap illustrating the separation of Purkinje-expressed genes into two clusters on the basis of the other genes they correlate with spatially. The *i*,*j*th entry is the number of genes found to overlap with both gene *i* and *j* in the Purkinje cluster (*4*). (**F**) For genes with significant expression (*p*<0.001) in the nodulus-uvula region (*4*), the fraction of reads localized to the nodulus/uvula and to the VI/VII boundary is shown. *Kctd12* and *Car7* did not pass the *p*-value cutoff, but are displayed as squares to demonstrate their location relative to *Aldoc*. (**G**) An *Aldoc* metagene, consisting of *Aldoc*, *Kctd12*, and *Car7* is shown in cyan. A *Cck* metagene, consisting of *Cck*, *Stmn4*, *Kcng4*, and *Atp6ap1l* is shown in red. (**H**) The *H2-D1* metagene consisting of *H2-D1*, *Cops7a*, and *Kmt2c* is shown in yellow. A *Hspb1* metagene, consisting of *Prkci* and *Hspb1*, is shown in blue. (**I**) As in (G), but only genes with significant expression both in the nodulus (p<0.05) and the VI/VII boundary (p<0.05) are shown. (**J**) The *Gnai1* metagene, consisting of *Gnai1*, *Nefh*, *Plcb4*, *Rgs8*, *Homer3*, *Scg2*, *Scn4b*, and *Gm14033* is shown in green, while the *B3galt5* metagene, consisting of *B3galt5*, *Gdf10*, *Tmem248*, *Mpped2*, and *Dpf3* is shown in magenta. (**K**) *Mybpc1*, a gene expressed in Bergmann glia, is shown in orange. (**L**) An Allen atlas image for *Mybpc1*. All scale bars show 250 µm; *Pcp4*, a ubiquitous marker for Purkinje cells is shown in gray in (C), (H) and (K).

Our algorithm also identified *Rasgrf1* as having significant nonrandom spatial distribution (*p*<0.001, N = 3 coronal cerebellar pucks) within the granule cell layer of the cerebellum (Fig. 3C, **cyan**), a pattern previously identified using ISH data (*10*) (Fig. 3D, **left**). We asked whether Slide-seq could discover any novel spatially patterned genes in the granular layer, which constitutes the main source of input to the cerebellar Purkinje cells. We divided the puck in Fig. 3C into anterior and ventral regions (**Fig. S6**) (*4*) and identified four genes (*Gprin3*, *Cemip*, *Syndig1l*, and *Mab21l2*) that showed strong and specific localization to the granule layer on the ventral side of the puck, especially lobules VIII through X (Fig. 3C, **yellow**). *Gprin3*, in particular, showed a distinct pattern, validated by the Allen Brain Atlas (Fig. 3D, **right**), which was largely the opposite of *Rasgrf1*.

The cerebellum is marked by parasagittal bands of gene expression in the Purkinje layer which are known to correlate with Purkinje cell physiology and projection targets (*11*–*14*). The most well-known of these genes is *Aldoc* (also known as the antigen of the Zebrin II antibody); several genes show similar or complementary parasagittal expression (*13*, *15*, *16*), but a systematic classification of banded gene patterns is lacking. We applied our spatial gene significance algorithm to the beads marked by NMFreg as Purkinje cells or Bergmann glia (the two resident cell types within the Purkinje layer) (*4*), identifying 669 candidate genes, including known markers of Purkinje banding, such as *Plcb4* and *Nefh*. Amongst these 669, we found 57 that correlate more with *Aldoc* than with *Plcb4*, a gene that is known to be expressed only in Zebrin II-negative bands (*17*), and 69 genes that correlate more with *Plcb4* than with *Aldoc* (Fig. 3E). Among the *Plcb4*-associated genes were 4 ATPases and 4 sodium channels: *Atp1a3*, *Atp1b1*, *Atp2b2*, *Atp6ap1l*, *Kcnab1*, *Kcnc3*, *Kcng4* and *Kcnma1*. Of these, *Kcng4* has previously been associated with increased firing rate in fast motor neurons (*18*), suggesting that its expression contributes to the faster spiking measured in Zebrin II-negative Purkinje neurons (*12*, *19*), while the calcium-dependent channel *Kcnma1* is known to regulate the timing of dendritic calcium burst spiking in Purkinje cells (*20*), suggesting that it contributes to differences in bursting activity previously observed between lobules III-V and X (*21*).

Beyond the classic parasagittal bands, previous studies have noted that the lobules of the cerebellum have distinct cognitive functions. In particular, lobules VI and VII have been associated primarily with cognitive tasks (*20*), whereas lobules IX and X have been associated with the vestibular system (*21*, *22*), so we asked whether there might also be differences in gene expression between these two regions. We therefore examined all genes with significantly enriched or depleted expression in each of these lobules (**Fig. S6**). Most of the genes thus identified displayed highly correlated expression in lobules IX/X and VI/VII (Fig. 3F), consistent with standard Zebrin II staining (Fig. 3G), but several, such as *Prkci, Prkcd* (*13*) and *Hspb1* (*23*), were restricted just to lobules IX and X. By contrast, other genes, which included *H2-D1*, *Cops7a*, and *Kmt2c*, displayed relatively uniform expression except in lobule X (Fig. 3H), confirming that lobules IX and X have their own independent program of gene expression.

We found other *Aldoc*-associated genes, exemplified by *B3galt5* (*4*, *23*), which showed exclusive expression in lobules IX/X and VI/VII (Fig. 3I), suggesting that these regions might share a third pattern of gene expression, despite the highly disparate cognitive roles typically associated with them. Consistent with the hypothesis of a third gene pattern, we also found several *Plcb4-*associated genes, including *Gnai1*, which had ubiquitous expression except in lobules IX/X and VI/VII (Fig. 3J). Remarkably, most of the genes that we found along with *B3galt5* are not Purkinje markers, suggesting that some spatial patterns in the cerebellum may extend to other cell types in addition to Purkinje cells. Indeed, the most prominent example we found is *Mybpc1*, a little-studied Bergmann cell marker that appears both in Slide-seq data (Fig. 3K) and in ISH data (Fig. 3L) to have a pattern of expression similar to *Aldoc, Kctd12*, and *Car7*. We thus conclude that the Purkinje cells, Bergmann glia, and granule cells of the cerebellum are all subdivided into many spatially segregated subpopulations. This finding is particularly surprising, since none of these distinctions was previously observed in single-cell sequencing studies (*7*, *24*). Although differences in Purkinje cell physiology across the cerebellum have been well-studied, we expect that future studies may reveal an as-yet-undiscovered, region-specific role for Bergmann glia as well.

Finally, to illustrate the utility of Slide-seq for studying biological responses to pathology, we applied it to a model of traumatic brain injury. Cortical injury was induced with an intracranial injection needle, and mice were sacrificed 2 hours, 3 days, or 2 weeks following injury. The puncture site was visualized in Slide-seq data by the presence of hemoglobin transcripts (at 2 hours, Fig. 4A), or transcripts of *Vim*, *Gfap*, and *Ctsd* (at 3 days and 2 weeks), and the regions of the puck thus identified matched the location of the injection site as determined by histology (Fig. 4B,C). Using the overlap algorithm developed for the sagittal cerebellum analysis, we identified all genes that correlated spatially with those transcripts. At the 2 hour timepoint, the only gene we found in this way was *Fos*, although we also found that rRNA correlated significantly with the hemoglobin genes (Fig. 4A) only at the injury site (**Fig. S7**).

**Figure 4:**
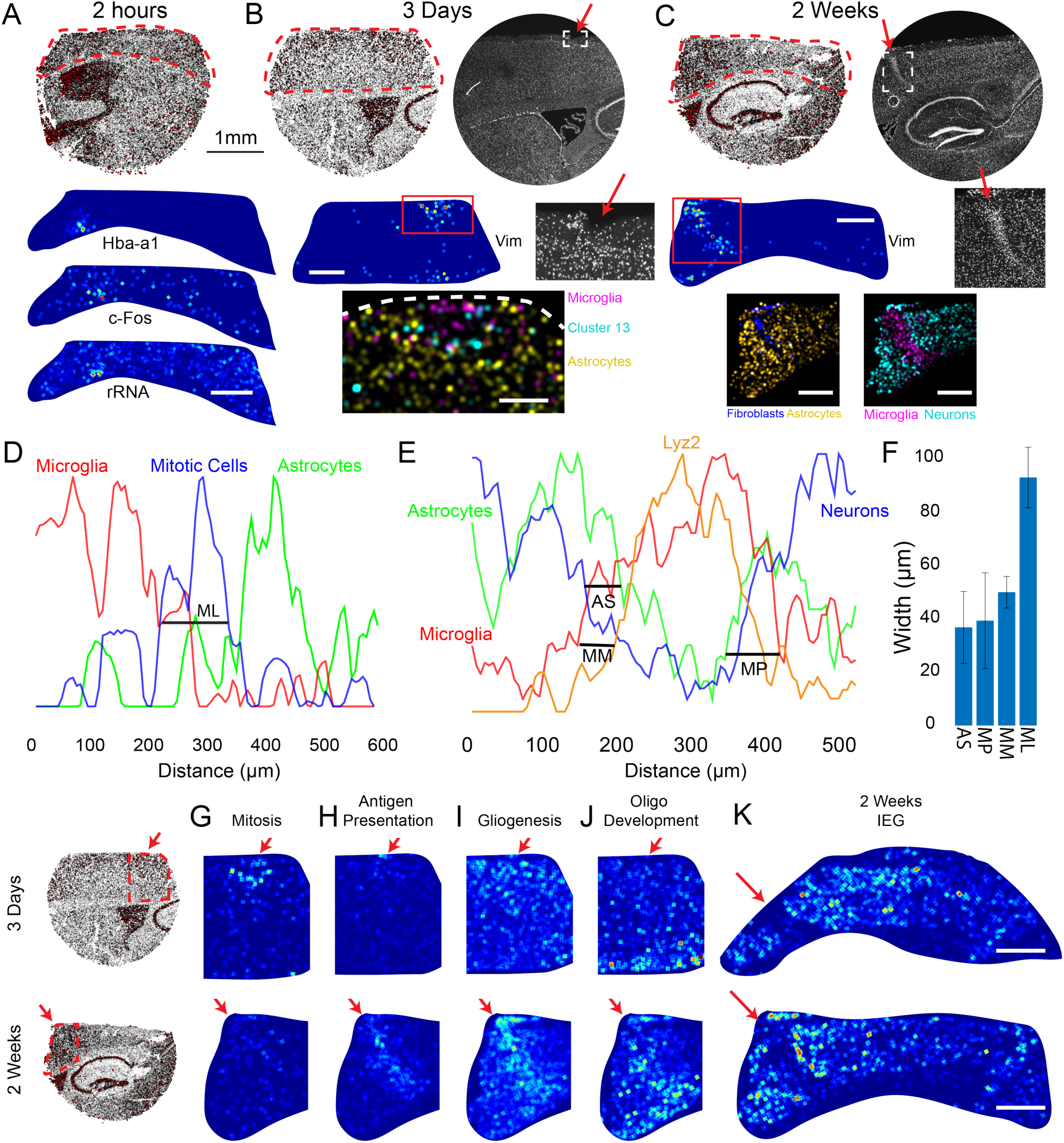
Slide-seq identifies local transcriptional responses to injury. (**A**) Top: All mapped beads plotted for a coronal slice from a mouse sacrificed 2 hours after injection. Circle radius is proportional to the number of transcripts per bead up to a maximum of 500 transcripts. Bottom: Three genes that mark the injection site. (**B**) As in (A), but for a mouse sacrificed 3 days after injection. Top right shows a DAPI image of an adjacent slice, with injury indicated by a red arrow and inset below. Middle right shows an expanded view of the injection site, with damage primarily in layer 1. Panels with black backgrounds show cell types as called by NMFReg using the hippocampal dataset, as density plots, scale bar: 250 µm. (*4*). Cluster 13 marks hippocampal neurogenesis, and is interpreted here as marking mitotic cells. (**C**) As in (B), for a mouse sacrificed 2 weeks following injection. Bottom, scale bar: 500 µm. (**D**) The spatial density profiles of microglia, mitotic cells, and astrocytes plotted for the 3-day puck in (B) (*4*). Lyz2 is taken to represent macrophages. (**E**) The spatial density profiles of microglia, astrocytes, and neurons for the 2-week puck in (C). (**F**) The average thickness of the features shown in (D) and (E) is shown, in microns. Error bars show standard error (N=6 for scar, N=6 for penetration, N=3 for mitosis layer). (**G-J**) Gene ontology-derived metagenes are plotted for the indicated transcriptional program, for two 3-day pucks (left) and two 2-week pucks (right). Red arrows indicate injection site. (**K**) The IEG metagene (see text) is plotted for two 2-week pucks. Circular images in (A), (B), and (C) refer to the 1mm scale bar in (A). All scale bars for images with blue backgrounds are 500 µm.

At the 3-day timepoint, in contrast to the 2-hour timepoint, we observed many genes at the injection site marking a robust astrocytic and immune response. Running NMFreg on the injection pucks, we observed a structured distribution of injury-associated cell types around the injection site at both the 3-day and 2-week timepoints. At the 3-day timepoint, we observed beads assigned to be microglia/macrophages localized to the lesion, bordered by a distinct layer of cells expressing mitosis-associated factors, followed by a layer of astrocytes (Fig. 4D), suggesting that mitosis occurs primarily in contact with the lesion itself. In contrast, at the 2-week timepoint, we observed microglia/macrophage-assigned beads filling the lesion, surrounded by consecutive layers of astrocytes and neurons, suggesting that mitosis had ceased by two weeks (Fig. 4E). Quantifying the thickness of these features (Fig. 4F), we found the mitotic layer at 3 days to be 92.4 µm ± 11.3 µm (mean ± sterr, N=3), suggesting several cell-widths of mitotic cells. By contrast, at 2 weeks, the astrocytic scar layer, defined as the distance between the half-maximum of the astrocyte layer and the half-maximum of the neuron layer, was 36.6 µm ± 13.4 µm (mean ± sterr, N=6), suggesting only one or two cell widths between the lesion and the nearest neurons. In addition, we also observed penetration of beads assigned to the microglia/macrophage through the astrocytic scar and into neuron-rich regions (39 µm ± 17.8 µm, N=6). Finally, we asked whether macrophages were localized differently from microglia. We plotted counts of the *Lyz2* gene, a specific marker for macrophages and granulocytes (*7*, *25*), and found that the distance between the half-maximum of the *Lyz2* distribution and the half-maximum of the microglia distribution was 49.7 µm ± 5.9 µm (mean ± sterr, N=6), suggesting that although microglia inhabit both the astrocytic layer of the scar and the lesion location itself, macrophages only inhabit the lesion. Thus, Slide-seq enables precise dissection of the spatial relationships amongst different cell-types, with resolution on the order of individual cells.

In order to investigate other changes in gene expression between the 3-day and 2-week timepoints, we ran spatial overlap analysis (*4*) to identify genes correlating with *Vim*, *Gfap*, and *Ctsd* at the 3 day timepoint and the 2 week timepoint. Applying gene ontology analysis to the set of genes correlated at one timepoint but not the other identified annotations relating to chromatid segregation, mitosis, and cell division at the 3 day timepoint (Fig. 4G), annotations relating to the immune response (Fig. 4H), gliogenesis (Fig. 4I) and oligodendrocyte development (Fig. 4J) at the 2 week timepoint. The degree to which oligodendrocyte progenitor cells (OPCs) differentiate into oligodendrocytes following a focal gray matter injury is controversial (*26*), however, we confirmed that both *Sox4* and *Sox10* localize to the region surrounding the injury, indicating the presence of immature oligodendrocytes (**Fig. S8**). Thus, we conclude that cell proliferation occurs on the order of days following injury, and that the period between 3 days and 2 weeks is critical for fate determination of newly born cells following traumatic brain injury.

We finally asked whether there might be other changes in gene expression in the surrounding tissue that are not immediately localized to the injection site. Remarkably, we noticed that several immediate early genes, including *Fos*, *Arc*, *Npas4*, and *Junb*, are upregulated in a large region around the injection site at both the 3-day and the 2-week timepoints (*27*–*29*). Running the overlap analysis on these genes at the 2-week timepoint revealed a number of other genes that also localized near the wound, including *Egr1*, *Egr4*, *Lmo4*, *Nr4a1*, *Slc16a13*, *Rgs4*, *Grin2b*, and *C1ql3* (Fig. 4K). This increase in gene expression decreased with distance away from the injection site and reached its half maximum 0.722 mm ± 0.191 mm away from the injection site (mean ± sterr, N=4 measurements). The neural specificity of these genes leads us to conclude that acute intracranial injury leads to alterations in the pattern of gene expression and possibly activity in nearby neurons for weeks following the injury.

Slide-seq enables the spatial analysis of gene expression in frozen tissue with high spatial resolution and easy scalability to large tissue volumes. The unbiased capture of transcripts enabled facile integration with large-scale scRNA-seq datasets, and discovery of novel spatially defined gene expression patterns in normal and diseased brain tissue. We anticipate that Slide-seq will play important roles in positioning molecularly defined cell types in complex tissues, and defining new molecular pathways involved in neuropathological states.

## Author Contributions

F.C. and E.Z.M. conceived of the idea and supervised the work. S.G.R. and R.R.S. developed the puck fabrication methods. S.G.R. developed the puck sequencing and base-calling pipeline. S.G.R. and E.M. made the sequenced pucks. R.R.S. developed the tissue processing and library preparation pipeline. S.G.R and R.R.S. performed experiments generating data for the manuscript, with assistance from C.A.M., L.M.C., and C.V. A.G. developed the NMFReg method and performed the associated analyses. J.W. performed some initial analyses to help motivate using NMF for spatial mapping of cell types. S.G.R. developed the analysis pipelines for identifying spatially non-random and significantly correlated genes. S.G.R., R.R.S., F.C., and E.Z.M. wrote the manuscript.

## Acknowledgments

We would like to acknowledge Edward Boyden for his support. We would like to thank Daniel Goodwin and Shahar Alon for useful discussions relating to the identification of rRNA in single-cell sequencing data. We would like to thank Jamie Marshall and Anna Greka for helpful discussions regarding kidney tissues. We would like to thank the McCarroll lab for the use of their Zamboni Drop-seq analysis pipeline. We acknowledge Jon Bloom for help with development of methods, and Velina Kozareva and Tushar Kamath for assistance with algorithm implementation.

This work was supported by an NIH New Innovator Award (DP2 AG058488-01), an NIH Early Independence Award (DP5, 1DP5OD024583), the Schmidt Fellows Program at the Broad Institute and the Stanley Center for Psychiatric Research. S.G.R. acknowledges funding through the Hertz Graduate Fellowship and the National Science Foundation Graduate Research Fellowship Program (award #1122374).

