## Supplementary Materials for "Slide-seq: A Scalable Technology for Measuring Genome-Wide Expression at High Spatial Resolution"

**Supplementary Materials:**
Materials and Methods
Figures S1-S8
Tables S1-S2

**Materials and Methods:**

*Beads:*
Beads were produced by the ChemGenes Corporation on one of two polystyrene supports (**Agilent 10 µm TOSOH Resin support and Custom Polystyrene supports from AM Biotech)**. Oligonucleotide synthesis was performed as described for Drop-seq(1). Beads were used with one of the two following sequences:

Sequence 1:
5'- PEG Linker- TTTT-PCT-
GCCGGTAATACGACTCACTATAGGGCTACACGACGCTCTTCCGATCTJJJJJTCTTCAGCGTTC
CCGAGAJJJJJJNNNNNNNNT30

Sequence 2:
5'- Linker-
TTTTTTTTCTACACGACGCTCTTCCGATCTJJJJJJJTCTTCAGCGTTCCCGAGAJJJJJJNNNNNN
NNT30

Here, PCT represents a photocleavable thymidine; J bases represent bases generated by split-pool barcoding, such that every oligo on a given bead has the same J bases; Ns represent bases generated by mixing, so every oligo on a given bead has different N bases; and T30 represents a string of 30 thymidines.

***Puck Preparation**:*
Pucks were prepared in batches of 20 to 30, which could then be stored indefinitely, dehydrated, at 4C. Glass coverslips (Bioptechs, 40-1313-0319) were attached to a miniature centrifuge (USA Scientific 2621-0016) using double sided tape. Subsequently, the coverslip was cleaned by spraying with 70% ethanol and wiping with lens paper (VWR 52846-007) A spray-on silicone formulation was then applled to the coverslip, the cover to the minifuge was closed, and the minifuge was turned on for 10 seconds to spin coat. The minifuge was then turned off and the cover opened, and liquid tape (Performix 24122000) was sprayed onto the coverslip. The minifuge was again closed and turned on for 10 seconds. The coverslip was then carefully removed from the minifuge, and a gasket (3 mm diameter Grace Biolabs, CW-50R-1.0) was placed on top of the coverslip and pressed down. Beads were diluted to a concentration of approximately 100,000 beads/uL in ultrapure water (Thermofisher, 10977015). Beads were pelleted and resuspended twice in ultrapure water, and 10uL of the resulting solution was pipetted into each position on the gasket. The coverslip-gasket filled with beads centrifuged at 40C, 850g for at least 30 minutes until the surface was dry.

The gasket was carefully removed from the dried coverslip. Gentle pipetting of water directly onto the pelleted bead pucks removed all beads except for those directly in contact with the liquid tape layer. Beads removed in this way could be stored at 4C for later use. As much water was removed from the resulting pucks as possible, and the pucks were left to dry.

***Puck Sequencing***:
Puck sequencing was performed using SOLiD chemistry in a Bioptechs FCS2 flowcell using a RP-1 peristaltic pump (Rainin), and a modular valve positioner (Hamilton). Flow rates between 1mL/min and 3mL/min were typical. Imaging was performed using a Nikon Eclipse Ti microscope with a Yokogawa CSU-W1 confocal scanner unit and an Andor Zyla 4.2 Plus camera. Images were acquired using a Nikon Plan Apo 10x/0.45 objective. After each ligation, we acquired four images: one using a 488nm laser and a 525/36 emission filter (MVI, 77074803); one using a 561nm laser and a 582/15 emission filter (MVI, FF01-582/15-25); one using a 561nm laser and a 624/40 emission filter (MVI, FF01-624/40-25); and one using a 647nm laser and a 705/72 emission filter (MVI, 77074329). The final stitched images were 6030 pixels by 6030 pixels.

Sequencing consisted of three steps: primer hybridization, ligation, and stripping. During primer hybridization, a primer was flowed into the flowcell at 5uM concentration in 4xSSC, and was allowed to sit for 20 minutes. Subsequently, the flowcell was washed in 3mL of SOLiD buffer F. Following instrument buffer wash, ligation mix was flowed into the chamber and allowed to sit for 20 minutes, before being flowed back into its original reservoir. Ligation mix was reused for ~10 ligations, before being replenished. Following ligation, the flowcell was washed again in instrument buffer, and we then flowed in 1.5mL of SOLiD buffer C, followed by 1.5mL of SOLiD buffer B, and repeated this step once again, to cleave the SOLiD sequencing oligo. We then washed the flowcell in instrument buffer and repeated the ligation step. After the second ligation step, 10mL of 80% formamide in water was flowed into the flowcell and left for 10 minutes. The flowcell was then washed in instrument buffer, and the process repeated with the next primer.

Ligation mix:
1x T4 DNA Ligase Buffer (Enzymatics)
6 U/uL T4 DNA Ligase (Rapid) (Enzymatics)
40x dilution of SOLiD SR-75 sequencing oligo.

***Image Processing and Basecalling***:
All image processing was performed using a custom-built processing suite in Matlab. Briefly, we acquired one image for puck after each ligation, and each image contained four color channels. First, color channels were co-registered to each other by thresholding the images and maximizing the cross-correlation between the thresholded images. Subsequently, for each puck, the images of each ligation were registered to the image of the first ligation using a SIFT-RANSAC image registration algorithm based on the VLFeat SIFT package in Matlab (*2*). Registered images were then base-called on a pixel-wise basis, as follows. First, the intensities in the Cy3 channel were multiplied by a factor of 0.5 and subtracted from the intensities in the TxR channel, which accounts for cross-talk between the channels which resulted from the excitation of TxR using the 561nm laser. Furthermore, for even-numbered ligations, the image of the previous ligation was multiplied by a factor of 0.4 and then subtracted on a channel-by-channel basis from the image of the even ligation. Each pixel was then called by intensity. For pucks made using the 180402 bead batch, we further enforced the expected base balance by including an additional step in which the intensities of the dimmest channels were progressively increased until each channel accounted for between 20% and 30% of the pixels in the center of the image.

Beads were subsequently identified from the base-called images as follows. Each pixel was assigned a number, the base 5 representation of which corresponds to the bases that were called at that pixel on each ligation. Every such number that occurred on at least 50 connected pixels in the image was determined to be a bead, represented by the centroid of the connected cluster.

SOLiD barcodes were then mapped to Illumina barcodes using a custom-built Matlab application that identifies the pairwise distance between all members of the two sets of barcodes. Pairs of SOLiD barcodes and Illumina barcodes were saved for further analysis if the two barcodes were separated by at most two edits, and if the mapping between the barcodes was unique, i.e. if there were no other barcodes at equal or lower edit distance to either barcode.

***Tissue Handling***:
Fresh frozen tissue was warmed to -20C in a cryostat (Leica CM3050S) for 20 minutes prior to handling. Tissue was then mounted onto a cutting block with OCT and sliced at a 5 degree cutting angle at 10 µm thickness. Pucks were then placed on the cutting stage and tissue was maneuvered onto the pucks. The tissue was then melted onto the puck by moving the puck off the stage and placing a finger on the bottom side of the glass. The puck was then removed from the cryostat and placed into a 1.5ml eppendorf tube. The sample library was then prepared as below. The remaining tissue was redeposited at -80 and stored for processing at a later date.

***Library preparation***:
*RNA Hybridization:*
Pucks in 1.5mL tubes were immersed in 200uL of hybridization buffer (6X SSC with 2U/uL Lucigen NxGen RNAse inhibitor) for 15 minutes at room temperature to allow for binding of the RNA to the oligos on the beads.

*First Strand Synthesis*
Subsequently, first strand synthesis was performed by incubating the pucks in RT solution for 1 hour at 42C.

RT solution:
- 75 ul H2O
- 40 ul Maxima 5x RT Buffer (Thermofisher, EP0751)
- 40 ul 20% Ficoll PM-400 (Sigma, F4375-10G)
- 20 ul 10 mM dNTPs (NEB N0477L)
- 5 ul RNase Inhibitor (Lucigen 30281)
- 10 ul 50 µM Template Switch Oligo (Qiagen #339414YCO0076714)
- 10 ul Maxima H- RTase (Thermofisher, EP0751)

*Tissue Digestion:*
200uL of 2X tissue digestion buffer was then added directly to the RT solution and the mixture was incubated at 37C for 40 minutes.

2X tissue digestion buffer:
- 200mM Tris-Cl pH 8
- 400mM NaCl
- 4% SDS
- 10mM EDTA
- 32U/mL Proteinase K (NEB P8107S)

*Library Amplification*
The solution was then pipetted up and down vigorously to remove beads from the surface, and the glass substrate was removed from the tube using forceps and discarded. 200ul of Wash Buffer was then added to the 400ul of tissue clearing and RT solution mix and the tube was then centrifuged for 3 minutes at 3000 RCF. The supernatant was then removed, the beads were resuspended in 200uL of Wash Buffer, and were centrifuged again. After repeating this procedure an additional 2 times, the beads were moved into a

200uL PCR strip tube, pelleted in a minifuge, and resuspended in 200uL of water. The beads were then pelleted and resuspended in library PCR mix and PCR was performed.

Wash Buffer:
  10 mM Tris pH 8.0
  1 mM EDTA
  0.01% Tween-20

Library PCR mix:
  23ul H20
  25ul of 2x Kapa Hifi Hotstart ready mix (Kapa Biosystems KK2601)
  1ul of 100 µm Truseq PCR handle primer (IDT)
  1ul of 100 µm SMART PCR primer (IDT)

PCR program:
  95 C 3 minutes
  4 cycles of:
    98 C 20 s
    65 C 45 s
    72 C 3 min
  9 cycles of:
    98 C 20 s
    67 C 20 s
    72 C 3 min
  Then:
  72 C 5 min
  4 C forever

### *PCR cleanup and Nextera Tagmentation*

The PCR product was then purified by adding 30ul of Ampure XP (Beckman Coulter A63880) beads to 50ul of PCR product. The samples were cleaned according to manufacturer's instructions and resuspended into 10ul of water. 1uL of the resulting sample was run on an Agilent Bioanalyzer High sensitivity DNA chip (Agilent 5067-4626) for quantification of the library. Then, 600 pg of PCR product was taken from the PCR product and prepared into Illumina sequencing libraries through tagmentation with Nextera XT kit (Illumina FC-131-1096). Tagmentation was performed according to manufacturer's instructions and the library was amplified with primers Truseq5 and N700 series barcoded index primers. The PCR program was as follows:

72°C for 3 minutes
95°C for 30 seconds
12 cycles of:
  95°C for 10 seconds
  55°C for 30 seconds
  72°C for 30 seconds
72°C for 5 minutes
Hold at 10°C

Samples were cleaned with AMPURE XP (Beckman Coulter A63880) beads in accordance with manufacturers instructions at a 0.6X bead/sample ratio (30uL of beads to 50uL of sample) and resuspended in 10uL of water. Library quantification was performed using the Bioanalyzer. Finally, the library concentration was normalized to 4nM for sequencing. Samples were sequenced on the Illumina NovaSeq S2 flowcell with 12 samples per run (6 samples per lane) with the read structure 42 bases Read 1, 8 bases i7 index read, 50 bases Read 2. Each puck received approximately 200M-400M reads, corresponding to 3,000-5,000 reads per bead.

***Calculation of Bead Packing***:

To estimate the packing fraction of the beads we imaged 10 pucks with 488nm light on the same microscope mentioned above after deposition onto the surface and prior to in situ sequencing. The signal was normalized to background and the image was binarized. The percent packing was reported as the fraction of the image occupied by the beads divided by the theoretical packing fraction of 0.9069 for dense packing of uniform spheres on a 2D surface. The mean and standard deviation of packing are reported in figure S1C.

***Clustering Analysis:***
For clustering of the pucks shown in Figure 1C, highly variable genes were identified by running FindVariableGenes() in the Seurat package in R, using a y.cutoff of 0.7 in liver, 0.6 in kidney and olfactory bulb, and 0.5 in hippocampus and cerebellum. For the hippocampus and cerebellum analyses, variable genes identified from the published datasets from these tissues were also included. Non-negative matrix factorization was performed using the NNLM package in R, on standardized, log-transformed values, with a k of 8 in liver and kidney, 6 in olfactory bulb, 13 in cerebellum, and 20 in hippocampus. The labeling each bead was of the largest factor loading from NMF after L2 normalization.

***Diffusion Analysis***:
An image of Slide-Seq bead signal density was generated through plotting the pixel intensity of each bead as a linear representation of the number of UMIs captured. The corresponding tissue slice was stained with DAPI and the width of the CA1 of each of the samples was measured through plotting a profile perpendicular of the feature with the intensity (either from UMI counts or DAPI signal) serving as the signal. The full width half maximum of the profile was then calculated for 10 such profiles across the CA1 for both Slide-seq and the serial tissue section as seen in Fig 1D.

***Image segmentation for cell estimates***:
For calculation of the total UMIs/ total cells estimated for each of the pucks images of DAPI stained serial sections were used. For tissues in which nuclei were easily separable at lower resolution we used 10x magnification and took a large stitched image. A region of the same area as the puck was then cropped from the image and used for segmentation/ counting of cells as described. For tissues where nuclei were not separable at 10X we took 10X images as well as large 60X images of dense "granular regions". At 60X cells were relatively easily separable and a joint count was made where a region of interest at 10X would be used for part of these tissues for nuceli counting. The number of nuclei in each region was proportioned appropriately and counts were combined. This strategy was used for the Granular cells of the cerebellum as well as the granular region of the olfactory bulb. Kidney, Liver, and Hippocampal sections were all segmented at 10X. Segmentation was performed in ImageJ by first scaling signal to the background and then binarizing the image. A 1.5 micron Gaussian Blur was applied across the image to help with oversegmentation and a watershed transform was applied to improve separation of adjacent nuclei. Nuclei were counted only if having a diameter of over 2um and less than 12um. The total number of UMIs from the puck was then divided by the number of nuclei obtained to generate the statistic total transcripts/ total cells.

***Comparison to Bulk sequencing***:

To compare the capture of Slide-seq to bulk RNAseq data we used a stranded mRNA Truseq kit (Illumina #20020594) to prepare stranded PolyA selection libraries from a dissected saggital mouse hippocampus. The libraries were sequenced and estimated TPMs were generated using Salmon (4). The ATPM measurement for Slide-seq data was generated by summing counts across all genes across a puck and then dividing each gene sum by (total UMI count/1million). This gives the ATPM which gives the estimated counts per gene per million counts. Since Slide-seq is a 3' enrichment technique we did not normalize by gene length. The per gene distribution for each of these values (bulk TPM and Slide-seq ATPM) was plotted and linear regression was performed giving an R = 0.89 showing good agreement between the two methods.

***Cell Type Deconvolution***:

For each bead, the contribution of each cell type to the RNA on that bead was computed using a custom method, implemented in Python, termed NMFreg (Non-Negative Matrix Factorization Regression). The method consists of two main steps: first, single-cell atlas data previously annotated with cell type identities (5) is used to derive a basis in reduced gene space (via NMF), and second, non-negative least squares (NNLS) regression is used to compute the loadings for each bead in that basis.

To perform NMF on the single-cell data, highly variable genes were first selected as in Saunders et. al (5), and NMF was performed using a specified number of factors (see below). Each of the factors (also termed basis vectors) was mapped to a unique atlas cell type, yielding interpretability of the basis. The cell type identity of a factor was established as the most frequent cell type of atlas cells with highest loading in this factor. Multiple factors contributing to the same cell type were aggregated by taking the L2 norm. Next, for each Slide-seq bead, we first computed the bead loadings in the basis using NNLS. The resulting matrix of loadings (with dimensions of the number of beads by the number of factors) suffers from the well-known non-identifiability native to NMF, and a scaling of these loadings is customary before further utilizing them. Therefore, each of the factor loadings were scaled to have unit variance equal to 1. Finally, the cell type of the bead was assigned based on the identity of the maximum factor loading.

For the implementation of NMFreg in Figures 2B and 2C, an adult mouse single-cell cerebellum dataset(5) was used to define the NMF basis, using a k (factor number) of 25. The published subcluster identities from this tissue were modified to remove clusters of cells outside of the Slide-seq-assayed anatomical region (e.g., cells from midbrain not seen on the puck) and to reduce the number of subpopulations, most especially from glial types. Specifically, all endothelial populations were merged together into one population, as were non-Bergmann astrocytes and oligodendrocytes. Interneurons not annotated as unipolar brush or Golgi (clusters 3-1, 3-2, 3-3, and 3-4)—which could not be assigned to a specific type in the published dataset—were also grouped together. Only Slide-seq beads with more than 15 unique genes containing counts were used in NNLS regression. For the implementation of NMFreg in Figure 2D, an adult hippocampus scRNA-seq dataset (5) was used in NMF setting k to 30 with 5 variable gene cutoff for bead inclusion. The first-level published cluster identities were used for bead assignment to cell types. For the implementation of NMFreg in Figures 3 and 4, the data was processed similar to figure 2D using published cerebellum (5) (Figure 3) or hippocampus(5) (Figure 4) datasets.

For the calculations in Fig. 2C and Fig. S4, we determined that a cell type was present on a bead if the L2 norm of the corresponding vector was at least 25% of the L2 norm of the vector of all factor loadings for that bead. Fig. S4 shows the numbers plotted in Fig. 2C as a function of this cutoff.

***Confidence Thresholding:***

For the computation in Fig. S4, we first performed NMFreg using only beads with at least 100 total transcripts. This decreases the number of beads called by 72.6% +/- 13.7% (mean+/-std over 7 cerebellar pucks). Interestingly, there was no relationship between the number of UMIs per bead and the confidence score of the bead. **(Fig. S4F)**.

The bead factor loadings returned by NMFreg are in general less pure than the factor loadings obtained for single-cell sequencing data, possibly reflecting both the sparsity of the Slide-Seq data and RNA contributions of other adjacent cell types. In order to determine whether a given bead could be confidently assigned to its highest contributing cell type, as in Fig S4, we computed a cell type specific single cell derived threshold. The threshold for a given cell type is the maximum loading of this celltype among all single cells assigned not to this cell type.

***Analysis of larger spatial feature sizes***:
The diameter of Slide-seq beads is 10 μm (original feature size). In an attempt to investigate the importance of the size of the features, we generated larger beads *in silico*, selecting artificial feature sizes of 20, 40, and 100 μm. Aggregate array features were performed by taking bead centroid locations obtained through SOLiD sequencing and forming a grid of defined size over the locations of the beads and aggregating beads within each region of the grid and treating the resulting data as a single bead.

***3D volume reconstruction of hippocampus***:
For Fig. 2D and E, beads assigned to hippocampus scRNA-seq clusters 4, 5, and 6 (CA fields and DG) from serial hippocampal Slide-seq sections were plotted in space. Sequential slices were roughly aligned by the density and shape of beads localized to hippocampal morphology. Alignments were refined with the ImageJ plugin TurboReg (*3*). Volumes were reconstructed in 3D by generating a 3D image stack with a sphere of diameter 12.5 μm with intensity proportional to number of UMIs centered on each bead centroid.

***Hippocampal Subtype Images***:
Metagenes for Fig 2F were identified from cell type specific atlas expression. The metagenes were:

| **Subtype** | **Metagene list** | **Atlas Cluster** |
|---|---|---|
| CA3/Hilum | Satb1, Scg2, Nap1l5, Fxyd6, C1ql3, Necab, Slc35f1, Nrsn1, Calb2 | 6 |
| CA2 | Adcy1, Pcp4, Rgs14 | 6 |
| Subiculum | Rxfp1, Fn1, Lxn, Nr4a2 | All beads |
| CA1 | Tenm3, Lypd1 | 5 |
| DG | Mef2c | 4 |
| Neurogenesis | Beads assigned to Atlas Cluster 13 (*5*) | 13 |

Metagenes were plotted via density plots (see below) on their corresponding Atlas clusters. Beads corresponding to Hippocampal atlas clusters 4, 5, and 6 (CA1, CA2/3, and DG) were displayed in light gray as a counterstain.

***Density Plots:***
For the density plot images in Fig. 2F, 3 and 4 (black backgrounds), we formed an image as follows. Each point P in the 6030x6030 images was assigned an intensity equal to the sum of the intensities of all beads with centroids lying within 44-pixel square centered on P. For 2B (black backgrounds) and 4B,C, each bead assigned to the indicated NMFreg cluster was assigned a unit intensity, while the intensity for each

bead in 3F was taken as the total number of transcripts belonging to genes in the indicated metagene. Finally, the images were passed through Gaussian filters with a standard deviation of 12 pixels.

For the images with blue backgrounds in Fig. 4, each bead was represented by a square of length 70 pixels on each side, with intensity equal to the total number of transcripts belonging to the set of genes indicated in the legend. Overlapping squares summed their intensities in the overlap region. For Fig. 4G-K, all the images within a given panel are normalized to the same values (i.e., the same colors represent the same values in all four images).

***Significant Gene Calling:***
Genes were identified as spatially non-random using a custom Matlab application (see Fig. S5). The set of pairwise Euclidean distances between all beads was calculated. For each cluster, genes were identified as candidates for the statistical significance analysis if they had an expression of at least 0.1 transcripts per bead within that cluster in the atlas reference dataset, or if the variance within that cluster in the atlas reference dataset was at least 0.01 transcripts squared and the ratio of the variance to the squared expression was at least 7.5 (an empirically determined value). Moreover, candidate genes for the statistical significance analysis were required to have at least one transcript on at least 15 beads.

To determine whether a transcript had a significantly non-random spatial distribution within a particular set of beads (for example, within the set of beads called as Purkinje neurons by NMFreg), we compared the distribution of pairwise distances between the beads expressing at least one count of that transcript to the distribution of pairwise distances between an identical number of beads, sampled randomly from all mapped beads on the puck with probability proportional to the total number of transcripts on the bead. (Rigorously, therefore, the spatial significance gene algorithm determines whether the spatial distribution of a particular transcript differs significantly from the spatial distribution of all transcripts.) Specifically, we generated 1000 such random samples, and for each sample calculated the distribution of pairwise distances. We then calculated the average distribution of pairwise distances, averages pairwise across all 1000 samples. Finally, we calculated the L1 norm between the distribution of pairwise distances for each of the 1000 random samples and the average distribution, and the L1 norm between the distribution of pairwise distances for the true sample of beads and the average distribution. We defined $p$ to be the fraction of random samples having distributions closer to the average distribution (under the L1 norm) than the true sample, and considered any genes with values $p<=0.005$. Due to the very high false-positive rate implied by this $p$ value (often as many as 4000 genes would pass the filters described above, implying ~20 false positives), statistically significant genes were identified as those that showed up consistently across biological replicates.

***Overlap Analysis:***
To identify genes that are significantly correlated or anticorrelated with other genes, we applied a custom Matlab algorithm. For simplicity of description, we consider the case of determining the genes that are correlated or anticorrelated with a particular gene, gene A. For each gene in the genome, we generated a "true" image in which each bead with at least one transcript of the gene was represented by a square of side length 100 pixels (~64 microns). Then, for each gene, we additionally generated 50 "random" images in which the same number of transcripts were redistributed across all beads with probability proportional to the number of reads per bead. We then calculated the elementwise inner product between the image of gene A and the 50 random images every other gene, and calculated the mean and standard deviation of the inner products. We then compared the mean and standard deviation to the inner products of the image for gene A with the true image of every other gene, obtaining a Z score for each gene. All genes with Z scores greater than 3 were deemed correlated, while those with Z scores less than 3 were deemed anticorrelated. Due to the high false discovery rate, we typically ran the algorithm with several test genes (equivalent to gene A), and across several pucks. We would then

consider genes only if they were correlated by the z=3 threshold with at least (for example) 2 genes on at least (for example) 2 pucks.

***Regional Significance Analysis:***
For several of the analyses in Fig. 3, we used the following procedure to determine whether the expression of a gene within a given region of the puck was significantly enriched or depleted. We divided Puck 180819_12 into 5 regions (Fig. S6): a dorsal region, a ventral region, a nodulus region, a nodulus-uvula region (consisting of the nodulus and the anterior uvula), and a VI-VII region, corresponding to the posterior side of lobule VI and the anterior side of lobule VII. The significance of a gene was then determined by a Fisher exact test performed on the contingency matrix [A, N-A; B, M-B], where A is the number of counts of the gene in the designated region, B is the number of counts outside of the designated region, N is the total number of counts of any gene in the designated region, and M is the total number of counts of any gene outside of the designated region. As in the case of the significant gene calling algorithm, this analysis could be performed on a subset of the beads on the puck. This procedure provides a list of genes with a significantly different pattern of expression within the designated region than outside of the designated region, regardless of whether the expression is elevated or depressed.

Compared to the significant gene calling algorithm above, this method is less general in that it is not able to find novel patterns of gene expression de novo, but it has greater statistical power.

***Identification of spatially variable genes in the cerebellar granular layer:***
We identified Gprin3 by finding all of the genes with significant expression ($p$<0.001) in the ventral part of puck 180819_12 compared to the dorsal part of the puck, for which more than 80% of the transcripts were in the ventral portion. This yielded three hemoglobin genes, *Th*, *Cemip, Gprin3, Mab21l2,* and *Syndig1l*. The three hemoglobin genes and *Th* were discarded because they were not expressed in granule cells.

***Identification of* Aldoc- *and* Plcb4-*associated genes in the cerebellar Purkinje layer:***
To identify the *Aldoc* and *Plcb4*-associated genes, we ran the significant gene calling algorithm on 14 cerebellar pucks (3 coronal, 11 sagittal), restricted to beads called as cluster 2 (Purkinje cells), cluster 7 (Bergmann glia), or the union of cluster 2 and 7 together. In this way, we identified 669 genes that were significant in at least one of pucks, which presumably includes many false positives. We then used the significance overlap gene to identify, for each of the 669 genes, the other genes in the set that correlate significantly with that gene in space on at least one puck. We then calculated, for each pair of genes in the set of 669, the magnitude of the overlap between the sets of correlating genes. To construct the matrix in Fig. 3E, we restricted that overlap matrix to the set of genes that have a larger overlap with *Aldoc* by at least 3 genes, or a larger overlap with Plcb4 by at least 3 genes. We used this procedure on the ground that the false positives in the set of 669 would for the most part be unlikely to have a greater overlap with *Aldoc* than with *Plcb4*.

For the purposes of displaying the matrix thus obtained in Fig. 3E, we first normalized the *i,j*th entry of the matrix by dividing as follows:

$$p_{i,j} \leftarrow \frac{p_{i,j}}{\sqrt{p_{i,i}p_{j,j}}}$$

We then divided each column of the resulting matrix by the sum of the column. Finally, because the resulting matrix was asymmetric, we summed the matrix and its transpose. For purposes of display, we then performed Ward clustering in Matlab and ordered them by cluster.

***Identification of* Hspb1 *pattern:***

To generate Fig. 3F, we included genes if they had significant expression in the nodulus-uvula region at p<0.001. We excluded *Ttr*, which was not expressed in Purkinje cells. For purposes of display, *Kctd12* and *Car7* were added to the graph as squares to help illustrate the clustering of *Aldoc*-like genes and *Cck*-like genes.

***Identification of* B3galt5 *pattern:***
To generate Fig. 3I, we included genes if they had significant expression in the nodulus at p<0.05 and significant expression in the VI-VII region at p<0.05.

***Distance Measurements for Injection Brains:***
The distance measurements in Fig. 4D,E were performed by plotting beads in each of cluster of interest with radius proportional to the number of transcripts per bead, with one transcript corresponding to a 25 pixel diameter and 500 corresponding to a 125 pixel diameter. This was done to ensure that beads with more transcripts were weighted more heavily when calculating the spatial profile of the cell types. We then drew boxes around the injection site and took line profiles (i.e., summed along one axis), to generate the profiles in 4D,E.

For measurements of the mitosis layer thickness, we took two measurements from one puck (Puck_180821_3, both sides of the injection site) and one measurement from a second puck (Puck_180819_19, the bottom side of the injection site). For measurements of the astrocyte scar thickness and the microglial penetration thickness, we took six measurements: two on each side of the scar from each of three pucks (Puck_180819_5, Puck_180819_6, and Puck_180819_7).

For the distance measurements in Fig. 4G, we plotted grayscale versions of the images in 4G, and took linecuts similar to those taken for the measurements in Fig. 4D,E. We took measurements from each side of the injection for puck 180819_7 (4G, bottom). We additionally took measurements from one side of the injection on pucks 180819_5 and 180819_6. We only used one side from those pucks on the grounds that the injection site was very close to the edge of the puck on one side.

Two of the three-day injection pucks, puck 180819_16 and 180819_18, were excluded from all distance measurements on the grounds that the tissue damage did not appear on the puck.

One two-week injection puck, puck 180819_8, was excluded from all distance measurements on the grounds that the tissue slice was more lateral than the other tissue slices. It showed neither enrichment of the immediate early genes around the injection site, nor a dip in astrocyte density in the middle of the scar, leading us to suspect that it was at the edge of the wound.

***Identification of rRNA in pucks:***
During analysis of the 2-hour injection pucks, we observed many counts of the Lars2 gene correlating with hemoglobins and cFos at the injection site. Upon investigation of the Lars2 gene, we found using RepeatMasker (http://www.repeatmasker.org/) that it has a rRNA-derived repeat in its 3’ UTR, leading us to hypothesize that the counts we observed of Lars2 might in fact be misaligned rRNA reads (*6*). Moreover, we found that the spatial distribution of Lars2 counts across the puck is highly correlated to the counts of rRNA, supporting this hypothesis. We thus used Lars2 as a proxy for rRNA expression in Fig. 4A.

### *Animal Handling Information:*
*Animal housing:*

Animals were group housed with a 12-hour light-dark schedule. All procedures involving animals at MIT were conducted in accordance with the US National Institutes of Health Guide for the Care and Use of Laboratory Animals under protocol number 1115-111-18 and approved by the Massachusetts Institute of Technology Committee on Animal Care. All procedures involving animals at the Broad Institute were conducted in accordance with the US National Institutes of Health Guide for the Care and Use of Laboratory Animals under protocol number 0120-09-16.

*Traumatic Brain Injury Model:*
Animals for the TBI model were anesthetized and processed according to a standard intracranial injection protocol. Specifically, mice were anesthetized using isofluorane and stereotactically restrained. Subsequently, an incision was made in the scalp and a hole was made in the skull using a dental drill. A Hamilton needle (32 gauge, 7803-04) was lowered to 2mm below the surface of the skull, and was then promptly retracted. The wound was closed using Vetbond, and the animal was allowed to recover. Mice were treated with Buprenorphine-SR and Meloxicam for analgesia. Mice were sacrificed by cardiac perfusion 2 hours, 3 days, or 2 weeks following the injection.

*Transcardial Perfusion:*
Animals were anesthetized by administration of isoflurane in a gas chamber flowing 3% isoflurane for 1 minute. Anesthesia was confirmed by checking for a negative tail pinch response. Animals were moved to a dissection tray and anesthesia was prolonged via a nose cone flowing 3% isoflurane for the duration of the procedure. Transcardial perfusions were performed with ice cold pH 7.4 HEPES buffer containing 110 mM NaCl, 10 mM HEPES, 25 mM glucose, 75 mM sucrose, 7.5 mM $MgCl_2$, and 2.5 mM KCl to remove blood from brain and other organs sampled. The appropriate organs were removed and frozen for 3 minutes in liquid nitrogen vapor and moved to -80C for long term storage.

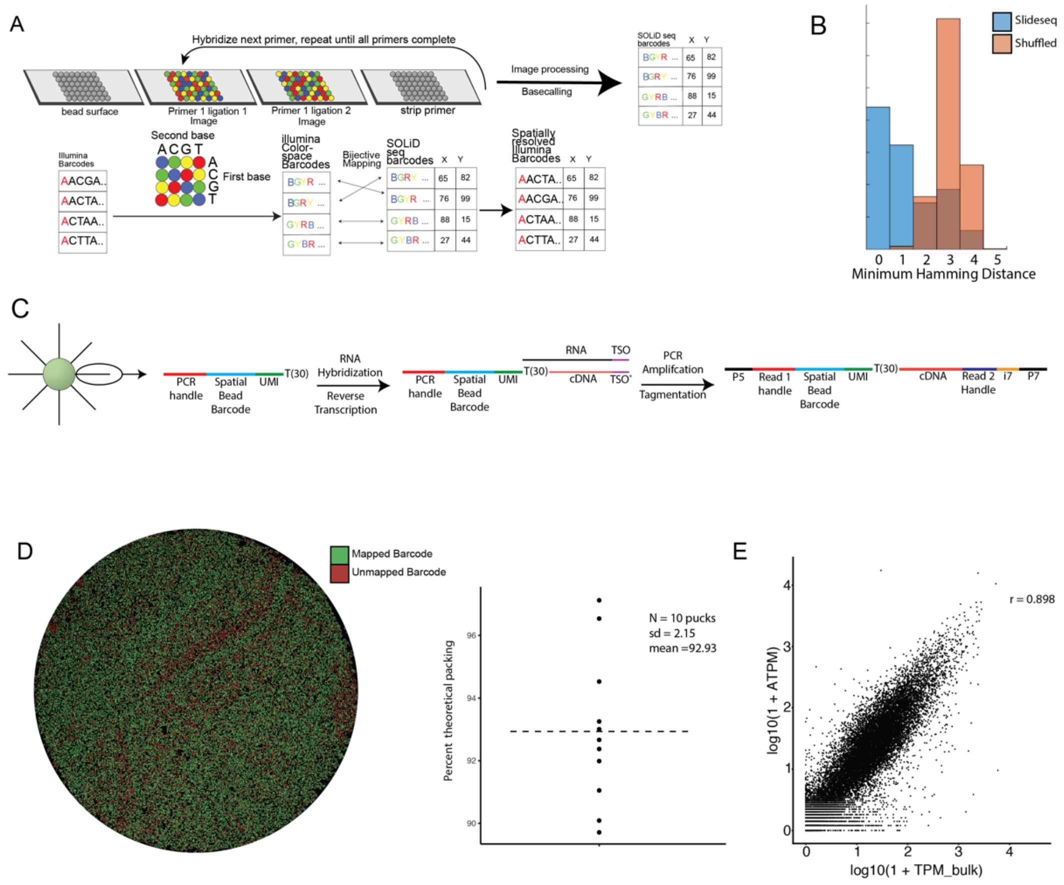

**Figure S1:** (**A**) Top: outline of the *in situ* sequencing and basecalling system established for generation of barcoded surfaces. Bottom: schema for mapping of Illumina barcodes to SOLiD barcodes. (**B**) Minimum hamming distance between Illumina colorspace-converted barcodes and barcodes from a puck sequenced *in situ* usign SOLiD chemistry (Blue, puck barcodes, Orange, shuffled puck barcodes). (**C)** Structure of the library at each stage of the preparation. (**D**) Barcode mapping across the puck. Beads colored green have a barcode bijectively matched between Illumina and SOLiD sequencing. Red beads are SOLiD-called barcodes not detected by Illumina sequencing. Beanplot shows the packing efficiency of the beads onto the surface. Mean ~ 93% the theoretical max of 90.7% for dense packing of uniform spheres(~85% bead occupancy). (**E**) Comparison of Slide-Seq count data to bulk RNAseq. X axis represents log10(1+TPM) of bulk data. Y axis represents log10(1+ATPM) of Slide-seq data (r = 0.89).

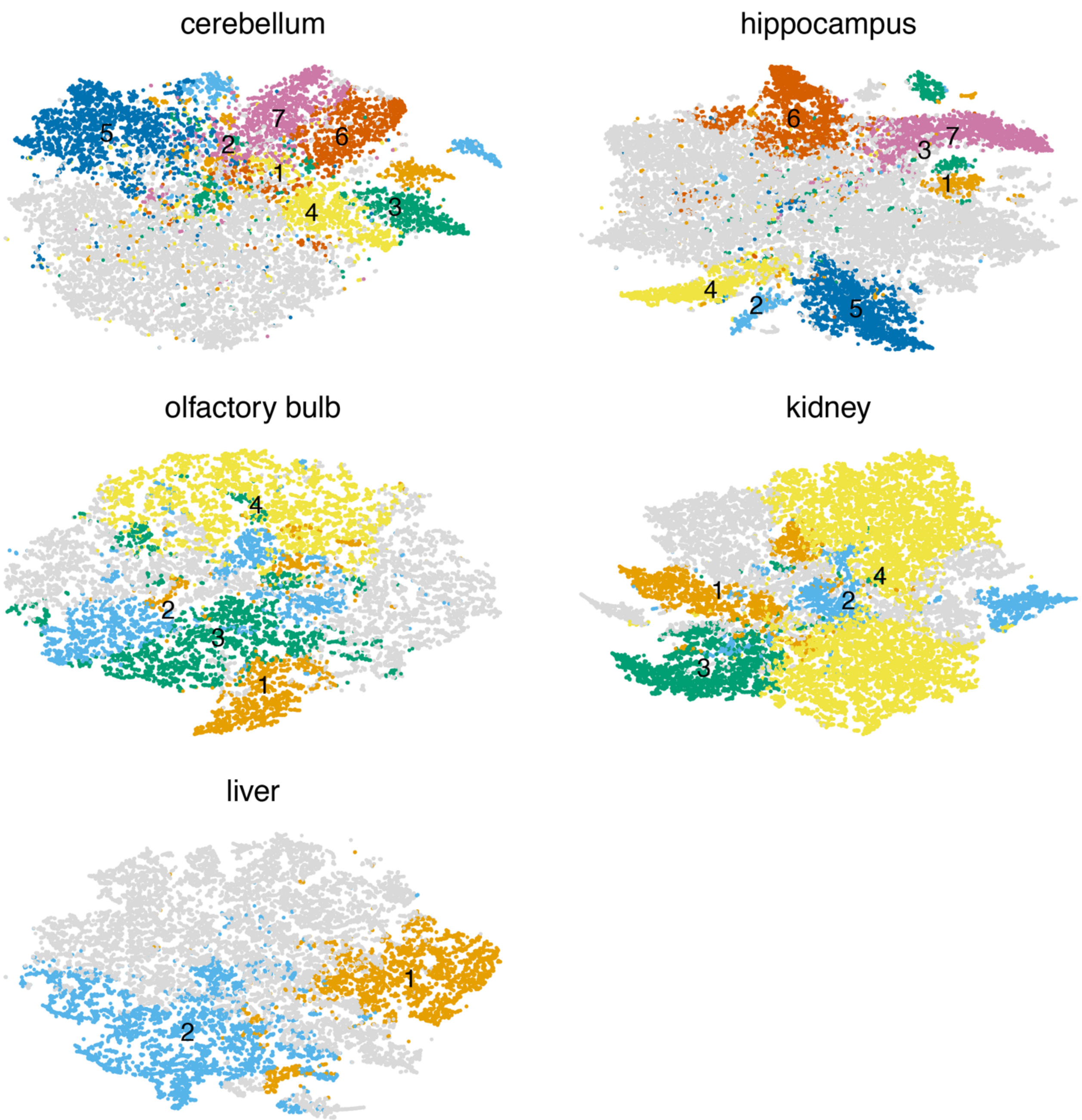


**Fig S2: Paired tSNE from Slide-seq data for various tissue types:** Shown are tSNE embeddings of the tissues assayed in Fig 1C. Coloring is consistent between the plots and highlights the ability of Slide-seq data to be unbiasedly clustered and spatially mapped to reveal underlying molecular features. Clusters not specifically highlighted in Fig 1C shown in gray.

Cluster identities were annotated as follows: Cerebellum: (1) Choroid plexus (2) Ependymal (3) cerebellar nucleus neurons (4) Cochlear nucleus (5) Oligodendrocyte (6) Purkinje cells (7) Bergmann glia. Hippocampus: (1) Fibroblast-like (2) ependymal (3) choroid (4) habenula (5) oligodendrocyte (6) CA1 neurons (7) dentate neurons. Olfactory bulb: (1) Glomerular layer (2) mitral layer (3) external plexiform layer (4) granule cell layer. Kidney: (1) Collecting tube (2) podocytes (3) Distal convoluted tubule (4) Proximal convoluted tubule. Liver: (1) Pericentral lobule layers (2) periportal lobule layers

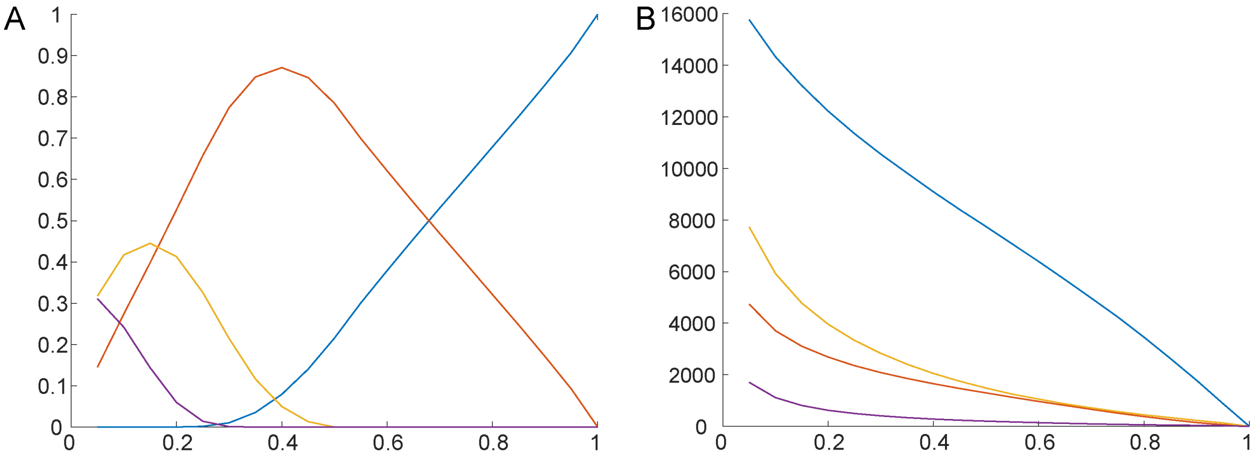


**Figure S3:** (**A**) A plot of the fraction of beads from cerebellar pucks analyzed in Fig. 2C, with zero cell types (blue), one cell type (red), two cell types (yellow), or three cell types (purple) as a function of the cutoff *C*. A cell type is defined to be present on a bead if the L2 norm of the vector of factor loadings mapping to that cell type is greater than or equal to *C* times the L2 norm of the vector of all factor loadings for that bead. For Fig. 2C, a cutoff of 0.25 was used. The plot shows mean across seven cerebellar pucks. (**B**) The mean number of beads representing granule cells (blue), Purkinje cells (red), other inhibitory neurons (yellow), and unipolar brush cells (purple) as a function of the cutoff *C*. The decrease in the number of each kind of cell is roughly linear for $C>0.4$, but is nonlinear for values of $C<0.4$, for which multiplets are possible.

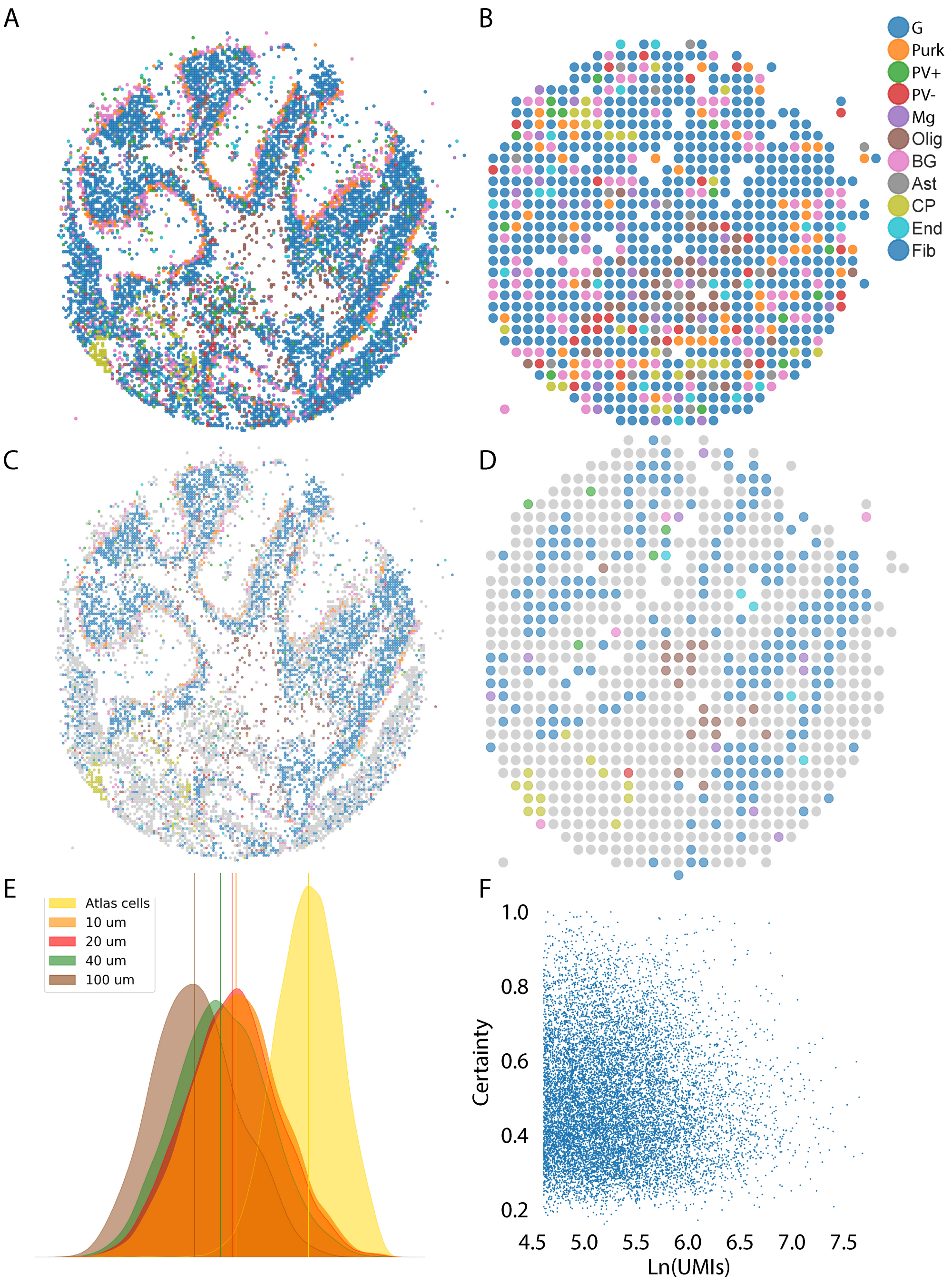


**Figure S4: Analysis of larger feature sizes, aggregated *in silico*.** (**A**) All beads were aggregated into 20 μm-diameter features and the resulting features were run through NMFreg. Beads are colored according to the cluster to which they were assigned, legend at right. G=Granule cells, Purk=Purkinje, PV+=Parvalbumin-positive interneuron, PV-=Parvalbumin-negative interneuron, Mg=Microglia, Olig=Oligodendrocytes, BG=Bergmann Glia, Ast=Astrocytes, CP=Choroid Plexus, End=Endothelium, Fib=Fibroblasts. (**B**) As in (A), but in this case beads were clumped into features with 100 μm diameter. Evidently, the structure of the tissue is largely lost. (**C**) Same as (A), but all features that fail to pass the confidence threshold are colored in gray. (**D**) As in (C), but for 100 μm features. Upon aggregating features into 100 μm diameter features, we retain the ability to identify choroid plexus, white matter, and granule cells, but no other cell types with confidence. (**E**) The distributions of L1 norms between the factor loading distributions and the uniform distribution are shown for atlas cells, the original Slide-Seq data (10 μm), 20 μm aggregated features, 40 μm aggregated features, and 100 μm aggregated features, showing the decrease in cell type purity as the feature size increases. (**F**) The number of UMIs (natural log) versus the confidence, defined as the L2 norm of the vector of factors mapping to the cell type as which the bead was called. There is no relationship between the number of UMIs and the bead confidence.

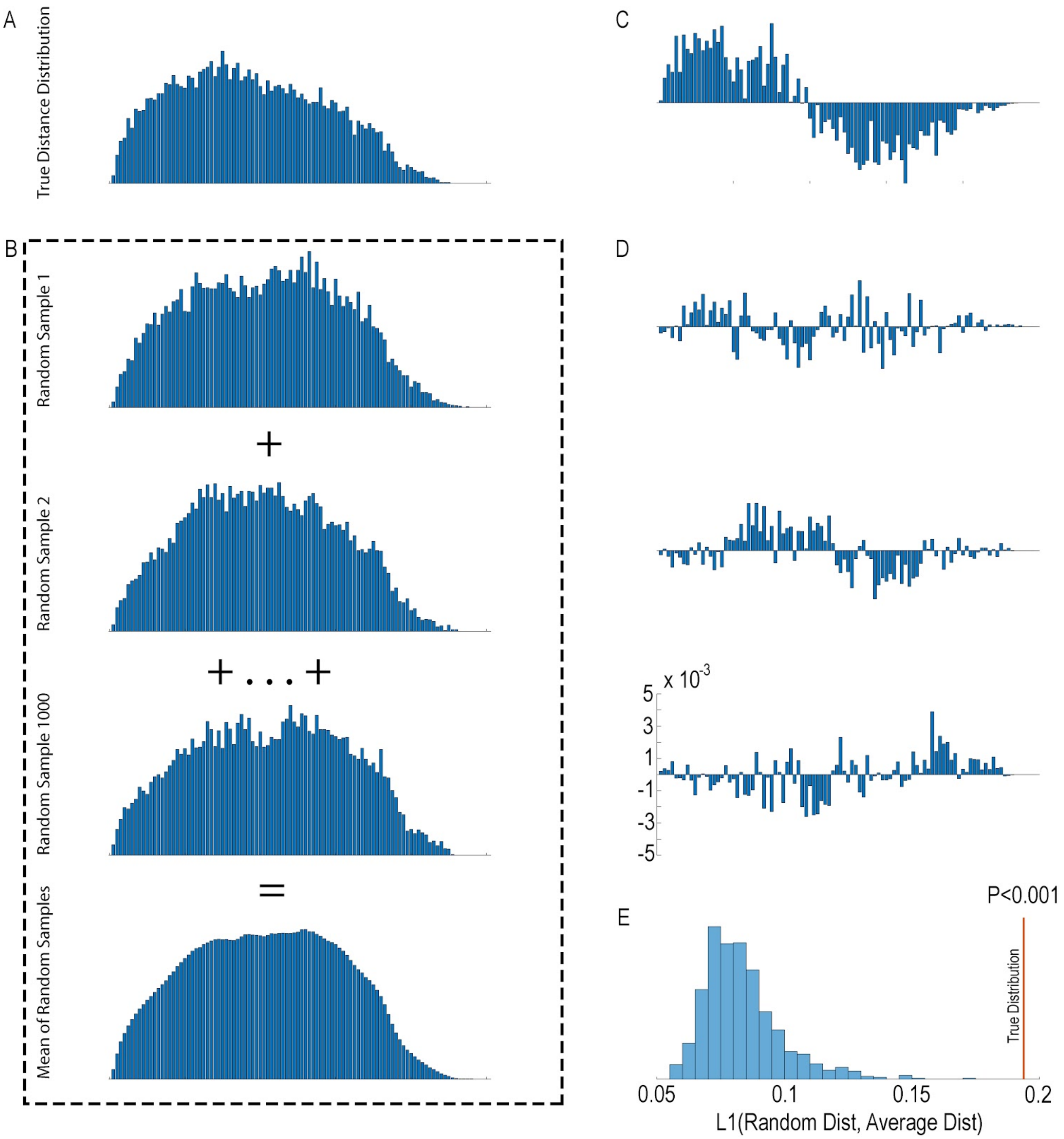


**Figure S5: Schematic of the significant gene calling algorithm**. The algorithm can be run on any specified subset of beads to identify genes with significant nonrandom distribution within that subset. All hisotgrams displayed here are calculated beads defined as granule cells on a coronal cerebellar puck (Fig. 3A). (A) For each gene of interest, we calculate the distribution of the Euclidean distances between all beads in the specified subset expressing at least one transcript of the gene, shown here for *Rasgrf1*. (B) We then randomly sample an equivalent number of beads from the subset with probability proportional to the number of reads per bead, without replacement. We perform this sampling 1000 times, and for each sample, calculate the distribution of pairwise Euclidean distances between the beads thus chosen. We take the elementwise mean of all 1000 samples to obtain the average distribution of pairwise distances across random samples. (C) We then take the elementwise difference between the distance distribution for the gene of interest and the average distribution, (D) as well as between the distance distribution for each of the random samples and the average distribution. (E) A histogram of the sum absolute values of the distributions shown in in (D), i.e., the L1 norm between distance distributions of the random samples and of the average sample. The L1 norm serves as our test statistic: if the gene of interest is distributed proportionally to the number of transcripts per bead, the L1 norm will be uniformly distributed. For *Rasgrf1*, the L1 norm of the true distribution is greater than the L1 norms of any of the random samples, so $p<0.001$. (Because there are only 1000 samples for reasons of computational complexity, the smallest observable p value is $p<0.001$).

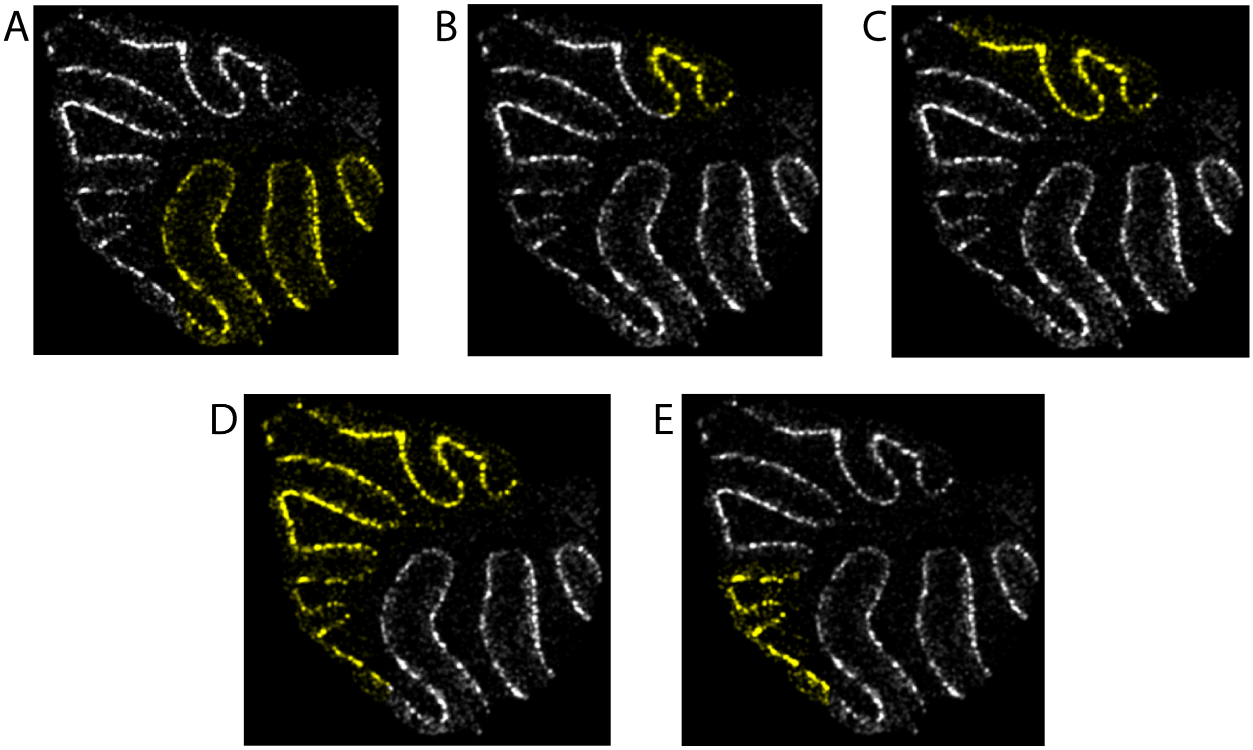


**Fig. S6:** Regions chosen for analysis in Fig. 3. Yellow indicates beads included in the region designation, while white indicates beads excluded from the region. A metagene consisting of Pcp4 and Pcp2 is plotted. (**A**) The dorsal region. (**B**) The nodulus region. (**C**) The nodulus-uvula region. (**D**) The ventral region. (**E**) The VIVII region.

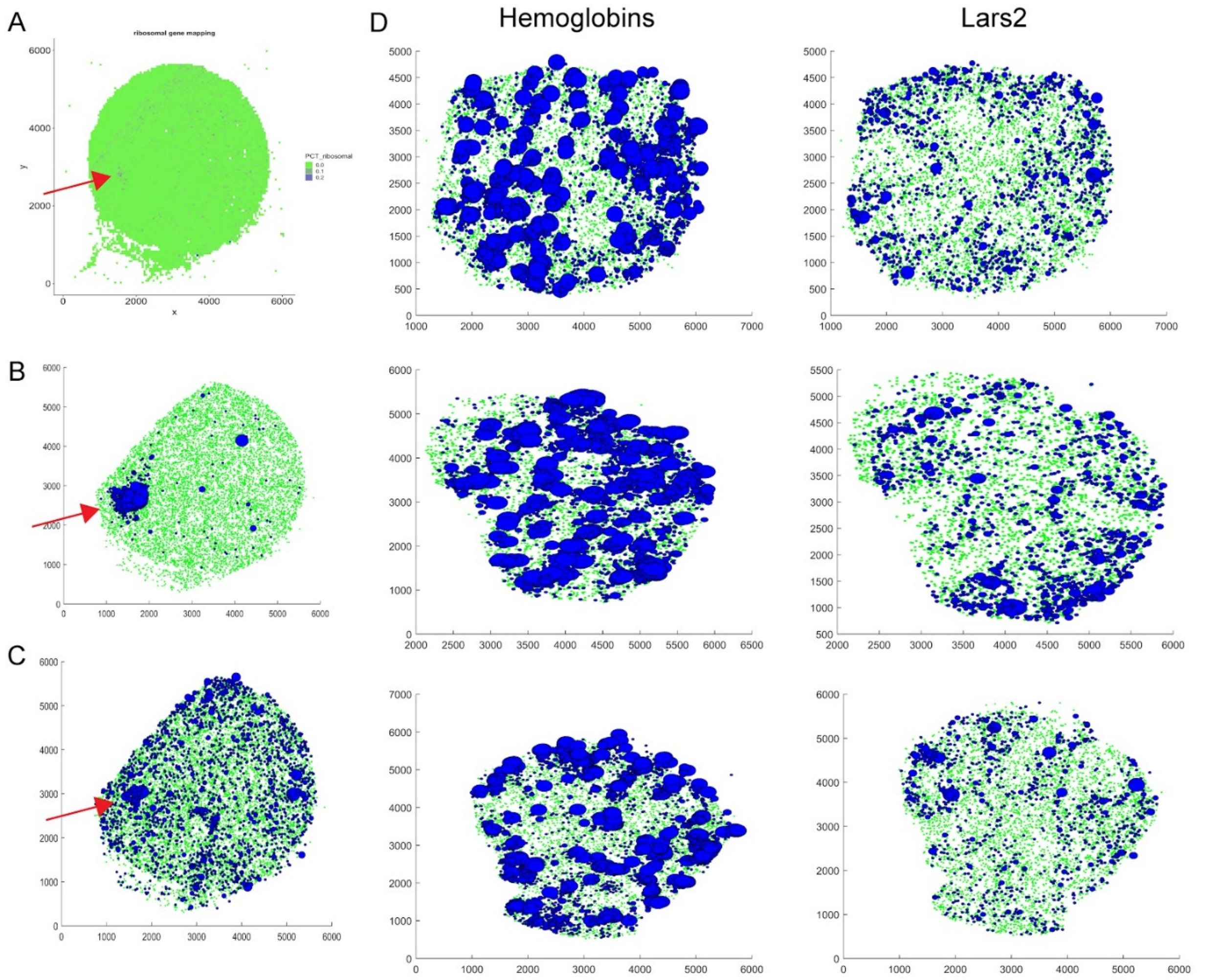


**Fig. S7:** (**A**) Plot showing the percentage of reads at each bead mapping to ribosomal RNA, prior to alignment, for the 180819_3 puck (same as in Fig. 4A) (**B**) Plot showing beads expressing hemoglobins. All beads expressing at least one transcript of Hba-a1, Hba-a2, Hbb-bs, or Hbb-bt are shown in blue, with radius proportional to the total number of hemoglobin transcripts. All other genes are shown in green. (**C**) As in B, but for Lars2 transcripts, which are believed to represent rRNA. (**D**) Three cerebellar (non-injected) pucks, showing hemoglobin transcripts (left) and Lars2 transcripts (right). The correlation between hemoglobin and Lars2 in B and C is in great excess over the correlations observed in D.

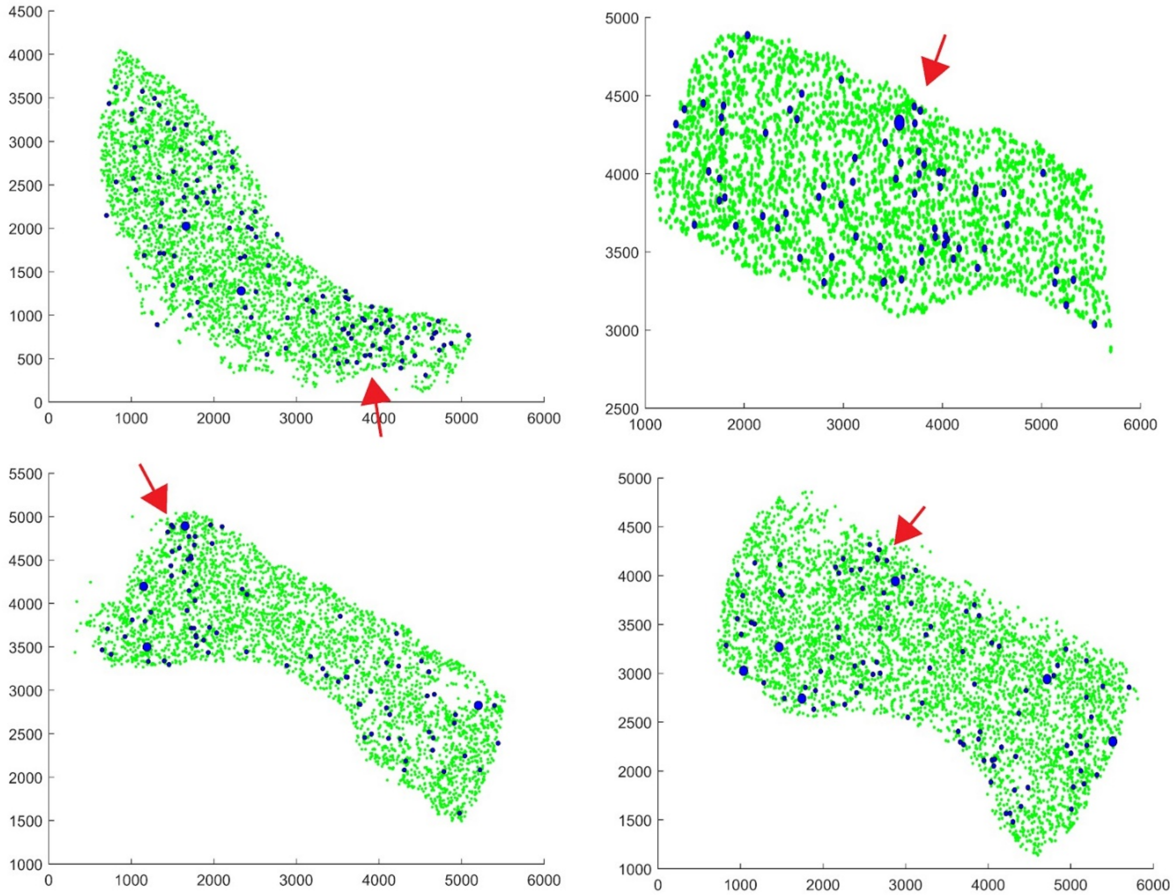


**Fig. S8:** Beads expressing *Sox4* and *Sox10* are shown in blue for four pucks from the 2 week injection timepoint. The radius of blue beads is proportional to the total counts of Sox4 and Sox10. The injection site is indicated with a red arrow.

**Supplementary Video 1:**
A 3D volume rendering of CA1, CA2/3 and dentate gyrus as shown in **Fig. 2**. Scale bars: 500 µm.

**Table S1:**
Oligonucleotides used in this study. Note r prior to base indicates RNA. + indicates LNA

| Name | Sequence |
|---|---|
| Truseq5 | AATGATACGGCGACCACCGAGATCTACACTCTTTCCCTACACGACGCTCTTCC GATCT |
| Smart PCR primer | AAGCAGTGGTATCAACGCAGAGT |
| Truseq_PCR_hand le | CTACACGACGCTCTTCCGATCT |
| Template Switch Oligo (TSO) | AAGCTGGTATCAACGCAGAGTGAATrG+GrG |

**Table S3:** Gene lists referenced throughout the paper, by figure.

| **Fig. 3** | |
|---|---|
| 669 Candidate Significant Genes | 1110001J03Rik 1700020I14Rik 1810037I17Rik 2210016L21Rik 2900093K20Rik AW047730 Abr Acin1 Actb Actr1a Actr3 Actr3b Acyp1 Adam11 Adam23 Add3 Aig1 Akap6 Akap9 Aldh5a1 Aldoc Alkbh7 Ank2 Ankrd12 Anks1b Ap1s1 Ap2a2 Aplp1 Apod Apoe App Appbp2 Ar Araf Arap2 Arfip2 Arhgap20 Arhgap5 Arl2 Arl4a Arpc4 Ascc1 Atp1a2 Atp1a3 Atp1b1 Atp1b2 Atp2b1 Atp2b2 Atp5c1 Atp5d Atp5h Atp5l Atp5o Atp6ap1l Atpif1 Atxn2 Atxn7l3b B230118H07Rik B2m Bag1 Baiap2 Bex2 Bhlhe41 Bloc1s6 Bola3 Brd7 Brd8 Brwd1 Bst2 Btbd17 Bzw1 Bzw2 Cacng2 Calb1 Calm2 Camk4 Capza2 Car2 Car7 Car8 Cbr1 Cbx6 Ccar1 Ccdc115 Ccdc50 Ccdc85b Ccdc88a Cck Cct6a Cd47 Cd63 Cd81 Cdc37l1 Cdc42ep4 Cdk5 Cdkal1 Cds2 Celf4 Cep126 Cept1 Cerk Cers4 Cggbp1 Chd9 Chga Chn1 Cisd3 Cit Ckap5 Clasp2 Cmtm5 Cnbp Cnot6l Cnp Col18a1 Commd7 Comt Copa Cops3 Cops4 Cops7a Cox14 Cox7a2l Cox8a Cpne2 Cpne9 Cr1l Creg1 Cript Cryab Csnk2a1 Cspg5 Cst3 Ctr9 Cttn Cttnbp2 Cux2 Cystm1 Cyth3 D10Jhu81e Dab1 Dagla Dap Dars Dbi Dclk1 Dcun1d5 Ddx1 Ddx42 Dgcr6 Dgkz Dnaja1 Dnajb2 Dner Dpm3 Dpp10 Dpysl2 Dstn Dtna Dync2li1 Ebf1 Echs1 Eci2 Ednrb Eif1ax Eif3a Eif3d Eif3f Eif4a1 Elmod1 Epb4.1l1 Epc1 Epha5 Ergic2 Erh Ermn Erp29 Etfa Evl Fabp3 Fabp5 Fabp7 Fam107a Fam174a Fam21 Fam98b Fbxl15 Fbxo3 Fbxo9 Fdps Fdx1 Fem1c Fgfr3 Fkbp1a Fkbp3 Fkbp8 Fth1 Fxyd7 Gabra1 Galnt11 Garnl3 Gas5 Gatm Gcsh Ggt7 Glul Gm14033 Gm27199 Gm5083 Gna13 Gnai1 Gnao1 Gnb2 Gng13 Gnl3l Golga4 Golph3 Got1 Gpatch11 Gpbp1 Gpm6b Gpr37l1 Gria1 Gria2 Gria4 Grid2 Grik1 Gsk3b Gstm1 Gtf2b Gtf2i Gucy1b3 Guk1 H2-D1 Hccs Hcfc1r1 Hdgf Hdlbp Hexa Hgsnat Higd2a Hint1 Hlf Hnrnpc Homer3 Hopx Hpcal1 Hprt Hsbp1 Hsd17b12 Hsf1 Hspa12a Hspa14 Hspa4l Hspe1 Hsph1 Hypk Icmt Id4 Ide Ifi27 Ifit3 Ifit3b Ifitm3 Ift57 Ilf2 Iltifb Ina Inpp5a Isca1 Itm2b Itm2c Itpr1 Jkamp Jrkl Kat6a Kcnab1 Kcnc1 Kcnc3 Kcnd2 Kcng4 Kcnma1 Kcnmb4 Kctd12 Khsrp Kif21a Kif3c Kif5c Kitl Klc1 Klhdc2 Kmt2c Krt25 Lamtor5 Lap3 Lars2 Ldha Lgals3bp Lhx1 Lhx1os Lin7a Lpcat4 Lpgat1 Lrrc49 Lsamp Luc7l3 Luzp2 Lztfl1 Macf1 Macrod1 Magoh Malat1 Map1a Map2k1 Map3k12 Mapk8ip2 Mapre2 Mapt March6 Mbnl2 Mbp Med8 Mef2a Meg3 Megf9 Mgst3 Mif Mipep Mir6236 Mkrn1 Mlec Mllt6 Mobp Morf4l2 Morn2 Mplkip Mrpl16 Mrpl35 Mrpl45 Mrps2 Mrps31 Msi1 Msi2 Msl3 Mt1 Mt2 Mt3 Mtdh Mtfmt Mtss1 Myo5a N6amt2 Nae1 Napg Nat8l Ncoa7 Ncor2 Ndufa11 Ndufa13 Ndufa2 Ndufa3 Ndufa4 Ndufa9 Ndufb2 Ndufb3 Ndufb4 Ndufb5 Ndufb8 Ndufb9 Ndufc1 Ndufc2 Ndufv1 Nefh Nefl Nefm Nnat Nomo1 Nop10 Npas3 Npc2 Npepps Nptx1 Npy Nr2c2 Nrsn1 Nrxn1 Nrxn2 Nsg1 Nt5c Ntrk2 Ntsr2 Nucks1 Oaz1 Oaz2 |

| | |
|---|---|
| | Ogfrl1 Olfm1 Omg Opa1 Opcml Opn3 Osbpl6 Ostc Pabpc1 Paip1 Pak1 Park7 Patz1 Pax6 Pbrm1 Pbx1 Pcdh17 Pcmt1 Pcp2 Pcp4 Pdcl Pde5a Pdhb Pdia3 Pdlim2 Pex13 Phip Pi4k2a Picalm Pigk Pigs Pisd Pitpnc1 Pja2 Plcb4 Plekhb1 Plekhb2 Plekhd1 Plp1 Pltp Pmm1 Pnn Pno1 Polb Polr2b Ppa1 Ppm1l Ppp1r11 Ppp1r12b Ppp1r17 Ppp2r2b Prdx1 Prdx3 Prdx5 Prdx6 Prex1 Prex2 Prkcd Prkcg Prkg1 Prkrir Prpf6 Psd2 Psma2 Psma3 Psmb10 Psmd8 Ptgds Ptpmt1 Ptpn11 Ptpn4 Ptprr Puf60 Pura Purb Pvalb Pxmp2 Qdpr Qk Rab24 Rabep1 Rabgap1l Rad23a Rad23b Ramp1 Ran Rasa2 Rasa3 Rbm5 Reep1 Rftn2 Rgs7bp Rgs8 Rims4 Riok2 Rit2 Rn18s-rs5 Rnf13 Rnf167 Rora Rpl14 Rpl18 Rpl34 Rpl38 Rpl41 Rps15a Rps21 Rps28 Rragc Rrp1 Rtfdc1 Rtn4 S100b Sac3d1 Saraf Scaf11 Sccpdh Scg2 Scn2a1 Scn4b Sdc3 Sdc4 Sdhc Senp2 Sep15 Sepp1 Sept11 Sept4 Sept7 Serbp1 Serinc1 Setd7 Sfxn4 Sigmar1 Slc13a5 Slc1a2 Slc1a3 Slc1a6 Slc24a2 Slc25a18 Slc25a39 Slc25a5 Slc33a1 Slc35a5 Slc38a1 Slc4a3 Slc4a4 Slc5a1 Smarca4 Smarcc1 Smpd1 Snap25 Snap47 Snapc3 Sncb Snhg11 Snrk Snrpn Snx24 Socs7 Sox9 Sparc Sparcl1 Spcs2 Sphkap Spock1 Spock2 Spred1 Srp9 Srsf2 Steap2 Stip1 Stk17b Stmn1 Stmn2 Stmn3 Stmn4 Strn3 Stt3b Stub1 Suclg1 Supt6 Sycp1 Syt2 Syt4 Syt7 Tardbp Tbc1d15 Tceb3 Tcf25 Tex261 Thy1 Thyn1 Timm10b Timm17b Tinf2 Tiprl Tln1 Tmed3 Tmed7 Tmeff2 Tmem11 Tmem158 Tmem167 Tmem184c Tmem255a Tmem47 Tmem50a Tmem50b Tmem64 Tmf1 Tmsb4x Tnik Tnrc6b Tomm22 Tomm40l Tpi1 Trf Trim2 Trp53bp1 Trpc3 Tsfm Tshz2 Tspan13 Tspyl4 Tst Ttc14 Ttc3 Ttl Ttyh1 Tuba1a Tubb2a Tubb2b Tubb4a Tubb5 Tulp4 U2af2 Ubap2l Ubb Ube2q1 Ube3a Ubfd1 Ubl5 Ubl7 Ublcp1 Uchl3 Ufc1 Upf2 Uqcr11 Uqcrb Uqcrh Usp14 Usp3 Usp33 Vcpip1 Vimp Vps26b Vps41 Wbp5 Wbscr22 Wdr33 Wdr7 Wwp1 Xrcc4 Ylpm1 Ywhah Zbtb20 Zcrb1 Zfc3h1 Zfp512 Zfp608 Zfp87 Zfr Zic1 Zmat2 |
| *Plcb4*-Associated Genes | Anks1b Atp1a3 Atp1b1 Atp2b2 Atp6ap1l Baiap2 Car8 Cck Cerk Chn1 Cops7a Garnl3 Gm14033 Gnai1 Golga4 Gria2 Grid2 H2-D1 Hdlbp Hnrnpc Homer3 Hpcal1 Hspa12a Icmt Ina Kcnab1 Kcnc3 Kcng4 Kcnma1 Kitl Kmt2c Lpgat1 Macf1 Mbnl2 Mef2a Msl3 Ndufb8 Nefh Nefm Nptx1 Pde5a Pja2 Plcb4 Pno1 Prdx5 Prkrir Qdpr Rabep1 Rgs7bp Rgs8 Riok2 Scg2 Scn4b Snhg11 Spock2 Stmn2 Stmn4 Strn3 Supt6 Thy1 Tmem50b Tmem64 Trim2 Tspan13 Ttc3 Vps26b Wdr7 Wwp1 Zbtb20 |
| *Aldoc*-Associated genes | Actb Aldoc Apoe Atp1a2 Atp1b2 Atp5l Atpif1 B230118H07Rik B2m Car7 Cd63 Cd81 Cdc42ep4 Cox14 Cpne9 Cst3 Dbi Dpm3 Dtna Ednrb Fam107a Fam98b Fth1 Glul Gpm6b Gpr37l1 Gria1 Gstm1 Hint1 Hopx Kctd12 Kif5c Mt1 Mt2 Mt3 Ndufa3 Ndufb4 Nomo1 Park7 Pigs Prdx6 Rpl34 Rpl38 Rpl41 S100b Sepp1 Sept4 Slc1a3 Sox9 Sparc Sparcl1 Suclg1 Tmem47 Tmsb4x Trf Tuba1a Zcrb1 |

| | |
|---|---|
| Genes with $p<0.001$ in the ventral part of puck 180819_12 compared to the dorsal part, and with greater than 80% of their counts in the ventral region. | Th Cemip Gprin3 Mab21l2 Syndig1l Hbb |
| Genes with $p<0.001$ in the nodulus-uvula region of puck 180819_12 (i.e. all genes appearing in Fig. 3E, except Kctd12 and Car7) | Aldoc Cacng4 Calm1 Calm2 Car8 Ccdc23 Cck Creg1 Cst3 Fabp7 Homer3 Hspb1 Idh3b Irs2 Malat1 Ngdn Plcb4 Prkcd Prkci Prpf31 Pvalb Rgs8 Slc1a6 Slc25a4 Sparc Stmn4 Ttr Uchl1 mt-Cytb mt-Rnr1 mt-Rnr2 |
| Genes with $p<0.05$ in the nodulus and $p<0.05$ in the VI/VII region of puck 180819_12 (i.e., all genes appearing in Fig. 3I). | Actb Aldoc B3galt5 Calm1 Car8 Cck Cdk5rap2 Chmp4b Cops3 Dbi Dpf3 Efr3a Eif5a Etfa Gad1 Gdf10 Gnai1 Gstm1 Homer3 Idh3g Itm2c Mpped2 Mybpc1 Nefh Nsg1 Plcb4 Ppp1r17 Pvalb Rabep1 Rgs8 Rims2 Rpl13 Sfxn1 Slc1a3 Sox9 Spock2 Timp4 Tmem248 Ttr Ufc1 Wbp2 Ywhah mt-Cytb mt-Rnr1 mt-Rnr2 |
| **Fig. 4** | |
| Genes correlating with Vim, Ctsd, and Gfap at the 3 day timepoint. | Camk2n1 Ctsd H2-T22 Hexb Lcn2 Lgals1 Mthfd1 Slc16a11 Pvrl3 Ttr Ctss Dbi Dhrs1 Fabp7 Gfap Mgp Mrps6 Mt2 Nupr1 Pea15a Pold4 Sdc4 Smc4 Trim30a Tspo Vim Vip B2m C1qc Fam124a Fth1 Gcnt2 Gzf1 Ifi27l2a Ifitm3 Myo6 Rpl22 Serpina3n Tnfaip8 Uimc1 Usp12 Vamp8 Xaf1 Ccdc115 Igfbp2 Igfbp7 Ubap2 Eif2ak2 2010111I01Rik Ccnd1 Cnot6l Efcab14 Gbp7 Maged2 Med17 Nfkbia Pabpc1 Rgs8 Rpl10a Smc2 Ugt8a Dclk3 Rnase4 Wnt7b Plp1 Trf Irf9 Rhoc S100a16 S100a6 Srgn Actb Apod Arpc1b Bcas1 Car2 Cldn11 Cnp Cplx3 Enpp2 Ermn Fam46a Gjc3 Grb14 Id1 Id3 Ifi27 Ifit1 Ifit3 Igfbp5 Irgm1 Isg15 Itgam Itm2b Lrp4 Lta4h Mag Mal Malat1 Mbp Mgst1 Mobp Mt1 Nipbl Psmb8 Pvrl1 Rhog Siglech Tppp3 Traf7 Fgfbp3 Creld2 Kcnip2 Msl3l2 Nfkb1 Nkd1 Stat3 Abca1 Aif1 Apbb1ip C1qa C1qb Calb1 Clic1 Cpne6 Crip1 Ctsb Cx3cr1 Cyba Dcps Fcer1g Ftl1 Fyb Gm14295 Grn H2-D1 H2-K1 Hba-a1 Hba-a2 Hbb-bs Hbb-bt Heg1 Hpgd Lcorl Lgals9 Ly86 Mpeg1 Msn Myl12a Myo1c Ncf1 Nes Nfe2l2 Nptxr Pkn1 Plek Ptbp3 Pycard Rn18s-rs5 S100a11 Slc44a2 Sparc Tle1 Tuba1c Tyrobp Uaca Vcan Xpnpep3 Igfn1 Lars2 Pdlim4 Prdx6 S100a13 Sept11 Sorbs1 Syt17 Tmem176b Aco1 Agtrap Bst2 Cald1 Cd63 Cd81 Chd1l Ctdspl Gbp3 Npas3 Ptpn13 Cd52 Ilk Pou2f2 Stat1 Ybx1 Ccnd2 Ctsz Nek6 |
| Genes correlating with Vim, Ctsd, and Gfap at the 2 week timepoint. | 1500015O10Rik 1700017B05Rik 1700047M11Rik 1810058I24Rik 2610015P09Rik 2810474O19Rik 3830403N18Rik 4632428N05Rik A2m AF251705 AW112010 Abca9 Abcb1a Abcd1 Abhd12 Abhd4 Abi3 Acads Acer3 Adam10 Adam17 Adamts1 Adamtsl4 Adap2 Add3 Adgre1 Aebp1 Afap1 Aff1 Agps Ahnak Ahr Aim2 Akap12 Akap13 Aldh16a1 Aldh1a1 Aldh2 Anapc7 Ang Angpt1 Ankrd13a Anxa2 Anxa3 Anxa4 Anxa5 Aplp1 Apobec1 Apobec3 Apoc1 Apoe Aqp4 Arap1 Arhgap17 Arhgap29 Arhgap30 Arhgdib Arrdc4 Arvcf As3mt Ascc2 Aspa Atf3 Atp1a2 Atp1b3 Atp6v0e Axl Bach1 Bcl2a1b Bfsp2 Bgn Bhlhe41 Bin1 Bin2 |

| | |
|---|---|
| | Blvrb Bmp2k Brd7 Bri3 Btg1 C3ar1 C4b Calr Capg Capns1 Carf Carhsp1 Casp8 Cav2 Ccdc13 Ccdc50 Ccdc74a Ccl3 Ccl4 Ccl5 Ccl6 Ccl9 Ccpg1os Cd14 Cd151 Cd164 Cd180 Cd302 Cd37 Cd44 Cd48 Cd53 Cd68 Cd74 Cd82 Cd83 Cd84 Cd86 Cd9 Cdc42ep4 Cdc42se1 Cdkn1c Cebpa Cebpg Cela1 Cenpb Cfh Cflar Cgnl1 Ch25h Chd4 Chst2 Clec5a Clec7a Clic4 Clmp Clu Cnn3 Cntrl Col12a1 Col1a1 Col1a2 Col27a1 Col3a1 Col4a2 Col5a1 Col6a1 Col9a3 Colec12 Colgalt1 Commd10 Coro1b Cotl1 Cpe Cped1 Cpne3 Cpq Cpt1a Cpxm1 Creg1 Crlf2 Crot Cryab Cryba4 Csf1 Csf1r Csf2rb Csrp1 Cst3 Cst7 Cstb Ctdsp2 Ctnna1 Ctnnb1 Ctsa Ctsc Ctsh Ctsk Ctsl Cttnbp2nl Cxcl14 Cxcl16 Cyb5r3 Cybb Cyfip1 Cyp4f14 Cyth3 Cyth4 Dab2 Dcn Ddah2 Ddr1 Diap2 Dio2 Dnase2a Dnm2 Dock1 Dock10 Dpp7 Dtx3l E130114P18Rik Edem1 Edn3 Ednrb Eef1a1 Eef1d Eef2 Ehd4 Eif3a Elf1 Elk3 Emid1 Eml4 Emp3 Endod1 Entpd1 Epas1 Epb4.1l2 Erbb2ip Erp44 Eya3 Ezr F11r Fabp5 Fam107a Fam114a1 Fam114a2 Fam46c Fblim1 Fbln1 Fbn1 Fcgr1 Fcgr2b Fcgr3 Fcho2 Fcrls Fermt3 Fgfr1 Fkbp7 Fli1 Flt1 Fmnl2 Fn1 Fnbp1 Fnip2 Foxc1 Foxo4 Frmd4a Fstl1 Fuca1 Fxyd1 Fxyd5 Gabarap Galnt10 Gatm Gbp2 Gcn1l1 Ghdc Gjb2 Gltp Glul Gm13139 Gm2a Gm973 Gna12 Gnai2 Gnb2l1 Gng12 Gng5 Gngt2 Gns Golim4 Golm1 Gpm6b Gpnmb Gpr183 Gpr34 Gpr37 Gpt Gpt2 Gpx1 Gsap Gsn Gstm1 Gstp1 Gucd1 Gusb Gyg H2-Aa H2-Ab1 H2-DMa H2-Eb1 H2-T23 H3f3b Hbegf Hdlbp Hes6 Hexa Hist1h1c Hist1h2bc Hk2 Hmha1 Hmox1 Hpgds Hrsp12 Hsd17b11 Hsd3b7 Hsp90b1 Hspb6 Hspb8 Hvcn1 Ifi30 Ifi35 Ifih1 Ifit2 Ifit3b Ifitm2 Ifnar1 Ifnar2 Ifngr1 Igbp1 Igf1 Igf2 Igfbp3 Ikbkb Il10rb Il21r Il33 Il6st Inpp5d Inppl1 Ipo8 Iqce Iqgap1 Irf8 Islr Itga6 Itgav Itgb1 Itgb3bp Itgb5 Itih5 Kcnj10 Kctd12 Kctd5 Kdm5a Kif5b Klf2 Klhl36 Klhl5 Klk6 Krcc1 Lactb Lactb2 Lair1 Lamb1 Lamb2 Lamc1 Lamp1 Lamp2 Lap3 Laptm4a Laptm5 Lat2 Lats2 Lcp1 Lgals3 Lgals3bp Lgmn Lhfpl2 Lilrb4 Lima1 Limch1 Lipa Lmo2 Lpar1 Lpcat1 Lpl Lrp10 Lsp1 Lsr Ltbr Ly6e Lyn Lyz2 Maf Mafb Magoh Magt1 Maml2 Man2b1 Map4k4 Marcks Matn4 Mcl1 Mdk Metap2 Mfap1b Mlc1 Mmp14 Mob1a Mob3b Mob3c Mog Mrpl52 Ms4a6c Msx1 Mt3 Mtdh Myh9 Mylip Myo18a Myo1f Myo9b Myoc Myof Naglu Nagpa Nbl1 Ncf2 Nckap1l Ncl Ndrg1 Neat1 Nek7 Nek9 Nfe2l3 Nfia Nhlrc3 Npc2 Npm1 Nrp1 Nrp2 Ntpcr Oard1 Oat Olfml1 Olfml3 Olig1 Opalin P2rx4 P2ry12 P2ry13 P4hb Pacsin3 Padi2 Palld Parp3 Pbrm1 Pbx3 Pbxip1 Pdcl Pde3b Pdgfra Pdia3 Pdlim2 Pdlim5 Pdpn Pex19 Pfn1 Phkg1 Phldb1 Phldb2 Pla2g15 Pla2g16 Pla2g7 Pld4 Plekhb1 Plekhf2 Plgrkt Plin2 Pllp Plod3 Pltp Plvap Plxdc2 Plxnb2 Pmp22 Ppap2b Ppfibp2 |

| | |
|---|---|
| | Ppp1r14b Ppp1r18 Prdx1 Prex1 Prex2 Prkcd Psap Psen1 Psme2b Ptgds Ptma Ptn Ptp4a2 Ptpn1 Ptpn18 Ptpn6 Ptprb Ptprc Ptprz1 Ptrf Ptrh1 Qdpr Qk Rab3il1 Rac2 Rad9a Ramp2 Rarres2 Rasgrp3 Rassf2 Rassf4 Rbms1 Rcan3 Rcn3 Reep3 Rel Renbp Rest Rgl2 Rgs10 Rgs5 Rhoa Rhoj Rhoq Rlbp1 Rnaset2a Rnaset2b Rnf130 Rnf141 Rnf213 Rock1 Rpl13a Rpl18 Rpl18a Rpl23 Rpl26 Rpl32 Rpl35a Rpl37 Rpl37a Rpl39 Rplp0 Rplp1 Rplp2 Rps10 Rps11 Rps14 Rps15a Rps20 Rps24 Rps26 Rps27l Rps3 Rps5 Rps9 Rras Rrbp1 Rtp4 Rufy1 Runx1 S100a1 S100a10 S100a4 S100b Sall1 Samd9l Samhd1 Samsn1 Sat1 Scamp2 Scara3 Scarb2 Scd1 Scd2 Scpep1 Scrg1 Sdc3 Selplg Sepp1 Sept10 Serinc3 Serpinb9 Serpine2 Serpinf1 Serpinh1 Sfrp4 Sgk1 Sgpl1 Sh3bp2 Sh3d19 Sh3glb1 Sh3pxd2a Sirpa Sirt2 Slain2 Slc11a1 Slc12a2 Slc14a1 Slc15a3 Slc16a1 Slc16a2 Slc1a2 Slc1a3 Slc25a10 Slc25a15 Slc25a18 Slc26a2 Slc29a3 Slc38a6 Slc39a1 Slc44a1 Slco2b1 Slfn5 Smarca5 Smg8 Smim3 Snhg18 Snx18 Snx5 Soat1 Sowahc Sox10 Sox12 Sox4 Sp1 Sp100 Sparcl1 Spata13 Spi1 Spp1 Spsb1 Sspn St3gal6 Stat2 Stat6 Stx2 Sulf1 Sult1a1 Susd6 Svil Tab2 Tagln2 Tap2 Tapbp Tcirg1 Tead1 Tec Tep1 Tgfb1 Tgfb2 Tgfb3 Tgfbr1 Tgfbr2 Tgif1 Thbd Thbs2 Thbs4 Timp1 Timp2 Timp3 Tlr3 Tm4sf1 Tmed10 Tmed3 Tmed5 Tmem119 Tmem123 Tmem150a Tmem170b Tmem176a Tmem18 Tmem47 Tmem86a Tmsb4x Tmtc2 Tnfaip8l2 Tnfrsf1a Tnni1 Toporsos Tpm2 Tpm3 Tpm4 Tpp1 Tpr Trem2 Trex1 Trim12a Trim25 Trim56 Trip11 Trp53i13 Tsc22d4 Tspan2 Tspan4 Ttc28 Ubald2 Ucp2 Unc93b1 Usp25 Ust Vamp5 Vasp Vat1 Vgll4 Vkorc1 Vps54 Vtn Wapal Wasf2 Wfdc17 Wipf1 Wls Wnk1 Wnt5a Wrn Wsb1 Wwtr1 Xlr Ybx3 Zbtb20 Zc3hav1 Zeb2 Zfhx3 Zfp36l1 Zfp703 Zic1 Zmiz1 Znfx1 Abca1 Actb Agtrap Aif1 Apbb1ip Apod Arpc1b B2m Bcas1 Bst2 C1qa C1qb C1qc Cald1 Car2 Ccnd1 Ccnd2 Cd52 Cd63 Cd81 Cldn11 Clic1 Cnp Crip1 Ctsb Ctsd Ctss Ctsz Cx3cr1 Cyba Dbi Dhrs1 Eif2ak2 Enpp2 Ermn Fabp7 Fam46a Fcer1g Fth1 Ftl1 Fyb Gbp3 Gcnt2 Gfap Grb14 Grn H2-D1 H2-K1 Hexb Id1 Id3 Ifi27 Ifi27l2a Ifit1 Ifit3 Ifitm3 Igfbp2 Igfbp5 Igfbp7 Itgam Itm2b Lcn2 Lgals1 Lgals9 Ly86 Mag Mal Malat1 Mbp Mgp Mgst1 Mobp Mpeg1 Mrps6 Msn Mt1 Mt2 Myl12a Myo6 Ncf1 Nek6 Nfe2l2 Nfkb1 Nfkbia Nupr1 Pabpc1 Pdlim4 Pea15a Plek Plp1 Pold4 Pou2f2 Prdx6 Psmb8 Ptbp3 Pycard Rhoc Rhog Rnase4 Rpl22 S100a11 S100a13 S100a16 S100a6 Sdc4 Serpina3n Siglech Sparc Stat1 Stat3 Tmem176b Trf Trim30a Tspo Ttr Tyrobp Uaca Vamp8 Vcan Vim Ybx1 |
